# Resistance-Mutant Rescue Establishes On-Target CHI3L1 Inhibition by CHI-DEL, a Brain-Penetrant Small Molecule with In Vivo Efficacy in Glioblastoma

**DOI:** 10.64898/2026.10.20.760326

**Authors:** Saurabh Upadhyay, Longfei Zhang, Marianna Teplova, Shumayila Khan, Moustafa Gabr

## Abstract

CHI3L1 (YKL-40) drives glioblastoma progression and immunosuppressive neuroimmune signaling, but as a non-enzymatic secreted glycoprotein it has lacked a genetic tool to confirm that small molecule effects are truly on-target. From a DNA-encoded small molecule library we identify **CHI-DEL**, a potent CHI3L1 ligand, and solve a 2.81 Å co-crystal structure (PDB 10JV) revealing a binding trajectory centered on Trp99. Structure-guided mutagenesis yields W99F, a compound-resistant CHI3L1 allele that retains native secretion, basal signaling, and thermal stability, unlike the destabilizing W99A control, establishing a clean chemical-genetic separation of biology from drug sensitivity. In isogenic knockout/rescue cells, wild-type rescue restores both phenotype and **CHI-DEL** sensitivity across invasion, proliferation, and a multi-node signaling program (pSTAT3 and NF-κB), while W99F rescue restores phenotype but confers resistance to **CHI-DEL** that is complete for some readouts and partial for others. This same epistasis was reproduced in human iPSC-derived microglia, where **CHI-DEL** reverses a GBM-conditioned-medium-induced phenotype in a CHI3L1- and Trp99-dependent manner. In vivo, **CHI-DEL** achieves brain and tumor exposure, engages its target and disrupts CHI3L1-CD44 signaling in a dose-dependent manner, remodels the tumor immune microenvironment, and confers a dose-dependent survival benefit that is specific to the active compound. Together, these data extend CHI3L1 small molecule inhibitor development beyond in vitro optimization, providing genetic proof of on-target pharmacology and a brain-penetrant therapeutic strategy for glioblastoma.

## Introduction

Glioblastoma (GBM) is the most common primary malignant brain tumor in adults and remains one of the most aggressive cancers of the central nervous system. Despite maximal surgical resection followed by radiotherapy and temozolomide, recurrence is nearly universal and long-term survival remains poor ^1^. The limited effectiveness of current therapies reflects diffuse tumor infiltration, extensive molecular and cellular heterogeneity, adaptive treatment resistance and a profoundly immunosuppressive tumor microenvironment. Targeted therapies and immune-checkpoint inhibitors have provided limited clinical benefit in GBM, emphasizing the need to identify therapeutically tractable factors that contribute to both tumor progression and microenvironmental remodeling ^2,3^.

Chitinase-3-like protein 1 (CHI3L1), also known as YKL-40, is a secreted glycoprotein implicated in inflammation, extracellular-matrix remodeling and cancer progression ^4^. CHI3L1 belongs to glycoside hydrolase family 18 but lacks chitinase activity because of substitutions within its catalytic motif ^5^. Nevertheless, it retains a conserved carbohydrate-binding cleft and interacts with glycans, proteins and cell-surface receptors to regulate processes associated with cell survival, proliferation, migration, angiogenesis and inflammatory responses. Elevated CHI3L1 expression has been observed in multiple malignancies and is frequently associated with aggressive disease, treatment resistance and unfavorable clinical outcomes ^6,7^.

CHI3L1 is particularly relevant to GBM. Increased CHI3L1 expression is associated with high-grade disease, the aggressive mesenchymal tumor state and poor patient survival ^8^. Within the GBM microenvironment, CHI3L1 is associated with malignant glioma cells, resident microglia and infiltrating macrophages, positioning it at the interface between tumor-intrinsic signaling and immune-microenvironmental regulation ^9^. Experimental studies have implicated CHI3L1 in GBM-cell proliferation, survival, migration, invasion, angiogenesis and resistance to therapy ^10^. CHI3L1-containing signaling networks have also been linked to the recruitment and functional polarization of tumor-associated myeloid cells, thereby supporting an immunosuppressive microenvironment ^4^. These findings suggest that CHI3L1 is not merely a prognostic biomarker but an active extracellular regulator of GBM pathogenesis and a potential therapeutic target.

Recent studies have established that CHI3L1 is pharmacologically ligandable. Small molecule binders have been identified through biochemical screening, structure-guided optimization and DNA-encoded library (DEL) approaches, and ligand-bound crystal structures have defined the central carbohydrate-binding cleft as a tractable recognition site ^11–15^. A persistent limitation across this body of work, however, is that on-target pharmacology has so far been inferred rather than proven. Cellular and in vivo activity has been attributed to CHI3L1 engagement on the basis of biochemical affinity and phenotypic correlation, without a means of testing, within an otherwise unperturbed biological system, whether a compound’s effect genuinely requires the binding site defined by structure. This gap is not incidental: CHI3L1 has no catalytic or enzymatic function, and unlike kinases and other enzyme targets, for which resistance mutations at a catalytic or active site provide a standard route to chemical-genetic validation, no comparable tool has existed for CHI3L1. Establishing genetic proof of on-target activity for a non-enzymatic, structurally defined small-molecule interaction has therefore remained an open problem, both for CHI3L1 specifically and for extracellular, non-catalytic targets more broadly.

DNA-encoded libraries provide access to large, chemically diverse compound collections while requiring only small quantities of purified target protein ^16^. However, enrichment during affinity selection does not independently establish direct or reproducible target engagement; off-DNA resynthesis followed by direct biophysical validation is therefore essential for distinguishing genuine binders from enrichment artifacts. Structural characterization of validated ligands can subsequently define their molecular recognition features and, in principle, provide a rational basis for engineering the kind of resistance mutation that has not previously been available for this target ^17^.

Here, we screened a DNA-encoded small molecule library against human CHI3L1 and resynthesized prioritized candidates without their DNA tags for direct binding analysis. Surface plasmon resonance identified **CHI-DEL** as a potent CHI3L1 binder. We subsequently determined the crystal structure of the CHI3L1-**CHI-DEL** complex, revealing occupancy of the central carbohydrate-binding cleft through a binding mode centered on Trp99. Using this structure, we designed a compound-resistant CHI3L1 point mutant that preserves native protein biology while selectively disrupting **CHI-DEL** engagement, providing the first chemical-genetic tool to test on-target pharmacology for this target. We used this tool to establish genetic proof that the cellular and human iPSC-microglial activity of **CHI-DEL** requires the structurally defined binding site, while in vivo activity was supported by tumor target engagement, dose dependence, and an inactive analog. These findings identify a structurally characterized small molecule ligand of CHI3L1, define a chemical-genetic strategy for validating on-target activity at a non-catalytic target, and provide a brain-penetrant lead for CHI3L1-directed therapy in GBM.

## Results

### A DNA-encoded small molecule screen identifies CHI-DEL as a selective CHI3L1 binder

A DNA-encoded library comprising approximately 1.4 billion compounds (AlphaMa DEL screening kit) was screened against recombinant human His-tagged CHI3L1 immobilized on Ni-NTA magnetic beads, enabling extensive coverage of chemical space under affinity-based selection conditions. The screening workflow (Fig. S1) consisted of three iterative rounds of affinity selection, each involving target immobilization, incubation with the pooled library, stringent washing to remove non-binding species, and heat-mediated elution of retained binders. A no-target control (NTC) selection was performed in parallel under identical conditions to identify nonspecifically enriched compounds. Recovered DNA was quantified by qPCR at each round (3.40 × 10^7^ and 1.38 × 10^6^ copies after rounds two and three, respectively, an approximately 24.6-fold decrease); this decrease reflected lower overall DNA recovery and, by itself, did not establish increased selection stringency or target-specific enrichment.

Following the third round, DNA barcodes from the enriched pool were PCR-amplified and subjected to next-generation sequencing, and reads were decoded into chemical identities and copy numbers using AlphaMa’s proprietary decoding pipeline. Enrichment was assessed by comparing copy numbers in the CHI3L1 selection relative to the NTC selection, visualized as two- and three-dimensional building-block enrichment plots (DataWarrior), in which each point represents a full library member positioned by its constituent building-block indices ^18^.

Of the candidates prioritized for off-DNA synthesis and direct biophysical validation, three are highlighted here on the basis of their enrichment magnitude: **Cpd A** and **Cpd B**, the two most highly enriched features identified against CHI3L1 (302.4-fold and 121.8-fold over the mean, respectively), and **CHI-DEL** (97.5-fold) (Fig. 1a,b). Despite lower DNA-tag enrichment than either comparator, only **CHI-DEL** produced reproducible, concentration-dependent binding to CHI3L1 by surface plasmon resonance (SPR); neither **Cpd A** nor **Cpd B** showed reproducible binding under the same conditions (Fig. S2 and S3). **CHI-DEL** bound CHI3L1 with a kinetically derived equilibrium dissociation constant (K_D_) of 192 ± 92 nM (mean ± SD). The representative kinetic experiment shown yielded k_a_ = 2.78 × 10^5^ M^-1^s^-1^, k_d_ = 7.90 × 10^-2^ s^-1^, and K_D_ of 284 nM (Fig. 1c). Single-concentration SPR profiling at 50 µM against a panel of 15 related and unrelated proteins, including the chitinase-like paralogs CHI3L2, CHIT1, and CHIA, and the CHI3L1-associated interactors RAGE, Galectin-3, IL13Rα2, and CD44, showed that the response of **CHI-DEL** to CHI3L1 (41.0 ± 0.7 RU) exceeded every counterscreen protein by at least 14-fold (Fig. 1d), establishing that this binding is specific to CHI3L1 rather than a general feature of related carbohydrate-binding proteins.

**Figure 1.**
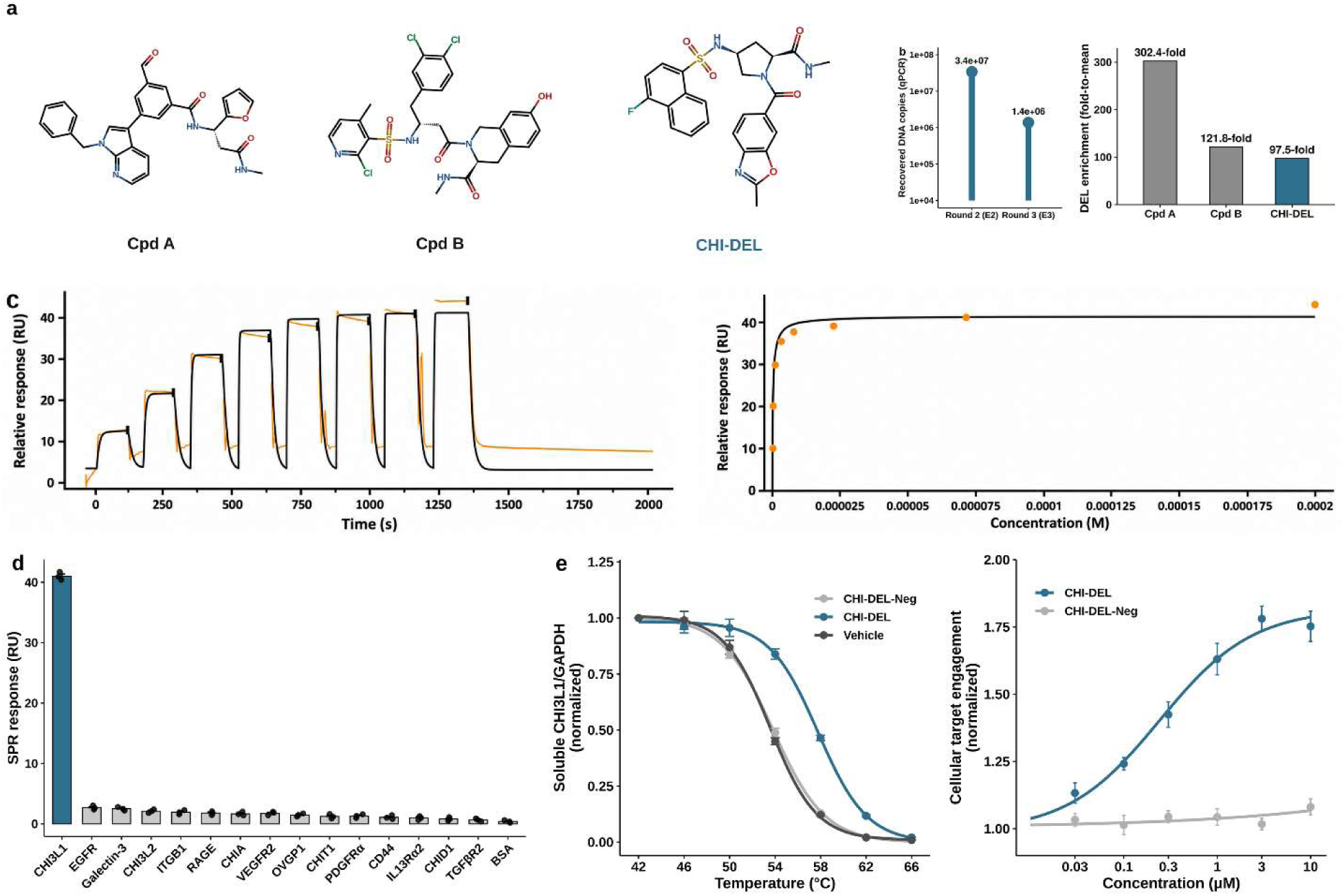
Discovery and validation of CHI-DEL, a selective small molecule CHI3L1 ligand. **(a)** Chemical structures of **CHI-DEL** and two comparator candidates (**Cpd A**, **Cpd B**) prioritized from a 1.4-billion-member DNA-encoded library screen against CHI3L1 for direct biophysical validation. **(b)** DNA-encoded library selection and enrichment. Left, recovered DNA copy number by qPCR after selection rounds two (E2) and three (E3), reflecting the change in DNA recovery across rounds. Right, DEL enrichment (fold-to-mean relative to the no-target control) for the three candidates shown in **(a)**. Despite 1.2- to 3.1-fold higher enrichment than **CHI-DEL**, neither **Cpd A** nor **Cpd B** produced reproducible SPR binding, demonstrating that enrichment magnitude alone does not predict target engagement. **(c)** Surface plasmon resonance analysis of **CHI-DEL** binding to immobilized CHI3L1. Left, representative single-cycle kinetic sensorgram (orange, experimental; black, 1:1 kinetic-model fit) across 0.091-200 µM, yielding K_D_ = 284 nM (k_a_ = 2.78 × 10^5^ M^-1^s^-1^, k_d_ = 7.90 × 10^-2^ s^-1^). Right, corresponding steady-state concentration-response. **(d)** Selectivity of **CHI-DEL** for CHI3L1 by single-concentration (50 µM) SPR counterscreen against 15 related and unrelated proteins, including the chitinase-like paralogs CHI3L2, CHIT1, and CHIA, and the CHI3L1-associated interactors RAGE, Galectin-3, IL13Rα2, and CD44. Individual replicate values (n = 3) are overlaid on the mean response for each protein. **(e)** Cellular target engagement in wild-type CHI3L1-expressing cells. Left, CETSA temperature-shift assay; **CHI-DEL** (1 µM) increases the apparent melting temperature of CHI3L1 relative to vehicle, whereas **CHI-DEL-Neg**, a structurally related analog sharing the same pyrrolidine core, does not. Right, isothermal dose-response CETSA; **CHI-DEL** engages CHI3L1 with EC_50_ = 0.33 ± 0.07 µM, while **CHI-DEL-Neg** is inactive across the same concentration range. Data are mean ± SEM, n = 3 biological replicates.

### CHI-DEL engages CHI3L1 in intact cells

To determine whether biochemical affinity of **CHI-DEL** translated into cellular target engagement, we used cellular thermal shift assays (CETSA). **CHI-DEL** (1 µM) increased the apparent melting temperature of soluble CHI3L1 relative to vehicle, whereas **CHI-DEL-Neg** (Fig. S4), a structurally related analog from the same DEL screen sharing the same pyrrolidine core, produced no shift (Fig. 1e, left). Isothermal dose-response CETSA confirmed this engagement was concentration-dependent (EC_50_ = 0.33 ± 0.07 µM), with no response to **CHI-DEL-Neg** across the same concentration range (Fig. 1e, right).

### Crystal structure reveals Trp99-centered recognition of CHI-DEL by CHI3L1

To define the structural basis of target engagement, we determined the crystal structure of a human CHI3L1-**CHI-DEL** complex at 2.81 Å resolution (PDB 10JV; Fig. 2a; Table S1). The asymmetric unit contains four CHI3L1 molecules, each bound to one **CHI-DEL** molecule. **CHI-DEL** adopts a similar orientation within the central carbohydrate-binding cleft in the four complexes and was modeled on the basis of the corresponding electron density (Fig. 2b). The ligand assumes an extended conformation within the cleft and is surrounded by residues lining both sides of the cavity (Fig. 2a,c). An electrostatic surface representation further illustrates the accommodation of **CHI-DEL** within the chemically heterogeneous central binding cleft, which contains negatively charged, positively charged and comparatively neutral surface regions (Fig. S5).

**Figure 2.**
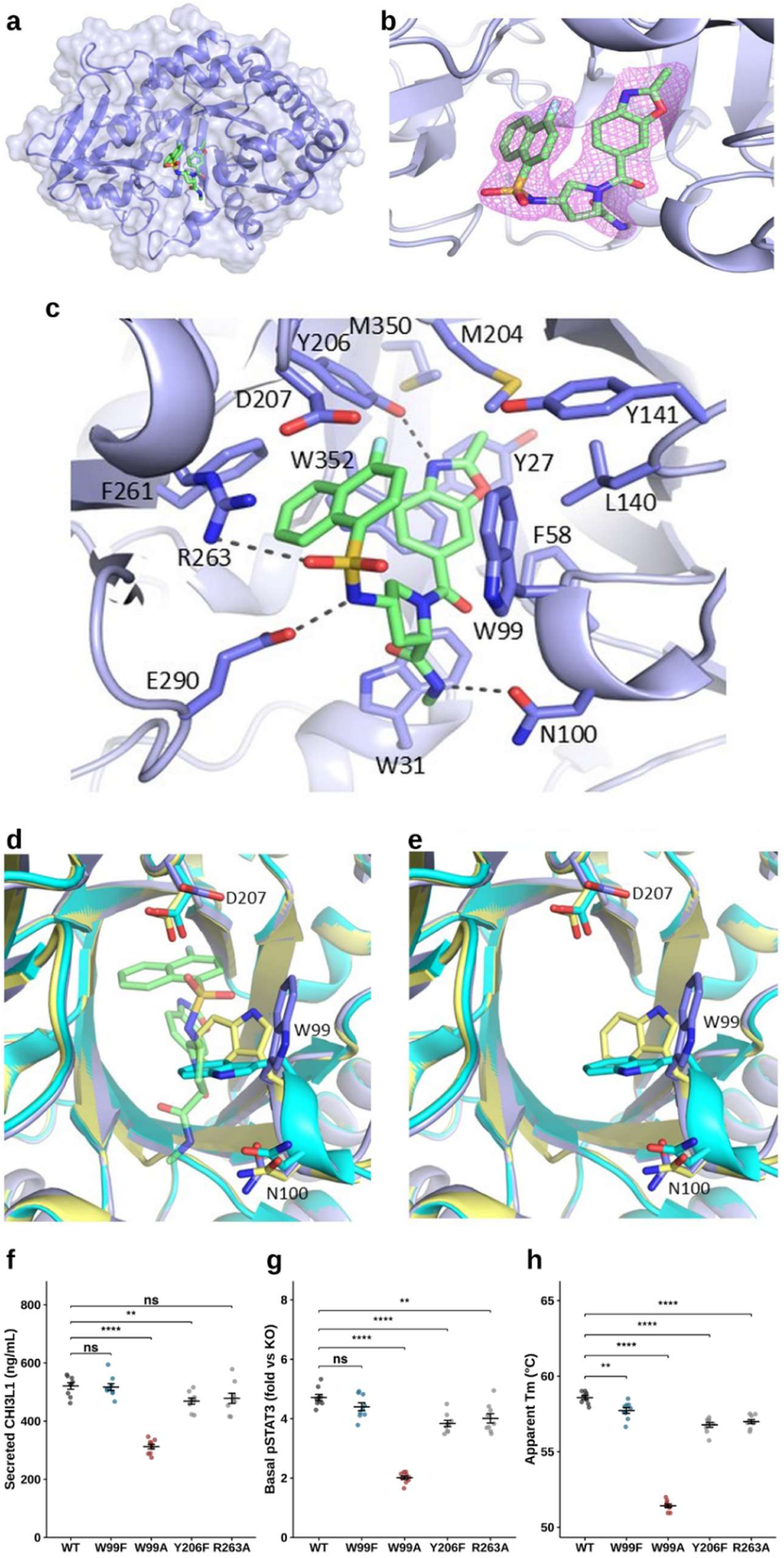
Crystal structure of the CHI3L1-CHI-DEL complex and structure-guided design of a resistance mutant. **(a)** Overall structure of CHI3L1 bound to **CHI-DEL** within the central carbohydrate-binding cleft (PDB 10JV). CHI3L1 is shown as a lavender cartoon with a semitransparent molecular surface, and **CHI-DEL** is shown as green sticks. **(b)** Close-up view of **CHI-DEL** within the 2Fo-Fc electron-density map contoured at 1.0σ, supporting the modeled orientation of the ligand within the binding cleft. **(c)** Molecular interactions between **CHI-DEL** and surrounding CHI3L1 residues. Key binding-site residues are shown as lavender sticks and labeled, and putative polar contacts are indicated by dashed lines. Oxygen, nitrogen, and sulfur atoms are colored red, blue, and yellow, respectively. **(d)** Superposition of the CHI3L1-**CHI-DEL** complex (lavender) with unliganded CHI3L1 (PDB 1HJX, light yellow) and inhibitor-bound CHI3L1 (PDB 8R41, cyan), highlighting **CHI-DEL** and the binding-site residues Trp99, Asn100, and Asp207. **CHI-DEL** is shown as light-green sticks. **(e)** The same superposition with **CHI-DEL** omitted to emphasize the different side-chain orientations adopted by Trp99, whereas Asn100 and Asp207 remain comparatively well aligned. **(f-h)** Biochemical qualification of WT CHI3L1 and four mutants (W99F, W99A, Y206F and R263A) prior to functional use: **(f)** secreted CHI3L1 by ELISA, **(g)** basal pSTAT3 signaling (fold over CHI3L1-KO baseline), and **(h)** apparent thermal melting temperature (T_m_) by differential scanning fluorimetry. W99F, Y206F, and R263A are statistically indistinguishable from WT on secretion and basal signaling and show only minor T_m_ shifts (≤1°C), whereas W99A shows substantially reduced secretion, blunted basal signaling, and a ∼7°C destabilization, identifying it as a loss-of-function allele unsuitable as a resistance mutant. Data are mean ± SEM; each point is one biological replicate (n = 9 per genotype). Brackets show comparisons to WT (Welch’s *t*-test, Benjamini-Hochberg corrected; ns, not significant; \**p* < 0.05, \*\**p* < 0.01, \*\*\**p* < 0.001, \*\*\*\**p* < 0.0001). Electrostatic surface representation of the binding cleft is provided in Fig. S5; crystallographic data collection and refinement statistics are provided in Table S1.

The ligand-binding cleft is lined by Tyr27, Trp31, Phe58, Trp99, Leu140, Tyr141, Met204, Tyr206, Phe261, Met350 and Trp352 (Fig. 2c). These predominantly aromatic and hydrophobic residues contact the nonpolar regions of **CHI-DEL**, consistent with extensive hydrophobic and van der Waals interactions. Geometrically plausible polar interactions involving Tyr206, Arg263 and Glu290 are consistently observed across the four crystallographically independent complexes. Asn100 and Asp207 lie adjacent to the ligand but are not assigned as direct polar-contact residues. Together, these interactions define the binding mode of **CHI-DEL** and provide a structural rationale consistent with its submicromolar affinity measured by SPR.

Superposition of the **CHI-DEL** complex with unliganded CHI3L1 (PDB 1HJX) and the previously reported inhibitor-bound structure (PDB 8R41) showed that the overall CHI3L1 fold is preserved upon **CHI-DEL** binding (Fig. 2d,e). The observed differences are concentrated within the ligand-recognition cleft. **CHI-DEL** follows a trajectory distinct from that of the inhibitor in PDB 8R41 and occupies the pocket through a different arrangement of hydrophobic and polar contacts. The most prominent local difference among the compared structures occurs at Trp99. This residue adopts different side-chain orientations in unliganded CHI3L1, the 8R41 inhibitor complex and the **CHI-DEL** complex (Fig. 2d,e), indicating conformational responsiveness at this position. In the **CHI-DEL** structure, the Trp99 side chain adopts an orientation that alters the local pocket geometry and enables complementary packing against the ligand scaffold. By contrast, Asp207 and Asn100 remain comparatively well aligned among the structures, suggesting that the most evident local difference in this comparison is associated with Trp99 rather than extensive rearrangement of the surrounding cleft.

This observation is consistent with the established contribution of Trp99 to CHI3L1 ligand recognition. In the GlcNAc₈-bound structure (PDB 1HJW), oligosaccharide binding is accompanied by a pronounced reorientation of the Trp99 side chain relative to unliganded CHI3L1, enabling stacking interactions with the carbohydrate ligand, together with displacement of Asn100 toward the binding cleft ^19^. Houston et al. also described conformational changes involving His209, Trp212 and Arg213, indicating that GlcNAc₈ binding induces a more extensive rearrangement of the recognition cleft than the predominantly Trp99-associated differences observed in our structural comparison. More recent ligand-bound structures have further demonstrated the contribution of Trp99-mediated CH-π interactions to small molecule recognition. Structure–activity analyses also indicated that engagement of the Trp99 region alone is insufficient for high-affinity binding and that complementary interactions within deeper regions of the cleft are required ^20^. These structural comparisons identify Trp99 as a ligand-responsive element through which CHI3L1 accommodates chemically distinct ligands. **CHI-DEL** stabilizes a distinct binding-site configuration involving Trp99 and reinforced by extensive contacts throughout the surrounding cavity. The structure defines the binding mode of **CHI-DEL** and identifies the Trp99-containing microenvironment as a potential focus for structure-guided optimization. Because **CHI-DEL** occupies the established carbohydrate-binding cleft, these findings also provide a basis for testing whether it competes with carbohydrate-dependent CHI3L1 interactions.

### Structure-guided design and qualification of a compound-resistant CHI3L1 mutant

The Trp99-centered recognition mode suggested that substitution at this position could selectively disrupt **CHI-DEL** binding. Because Trp99 also participates in the native ligand-recognition chemistry of CHI3L1, we reasoned that an aggressive substitution risked destabilizing the protein, confounding interpretation of any downstream resistance phenotype. We therefore designed a conservative substitution, W99F, preserving the aromatic ring while removing the indole nitrogen, alongside a more disruptive control substitution, W99A, and two additional pocket-adjacent mutants, Y206F and R263A.

Before using any functional assay, all four mutants were qualified against wild-type CHI3L1 across three independent readouts: secretion, basal pSTAT3 signaling, and thermal stability (Fig. 2f-h). Full melt curves are shown in Fig. S6a. W99F, Y206F, and R263A were statistically indistinguishable from WT in secretion and basal signaling, with thermal stability shifts of ≤1°C, an order of magnitude smaller than the ∼7°C destabilization observed for W99A, which also showed significantly reduced secretion and blunted basal signaling. W99F showed a modest reduction in intracellular CHI3L1 level relative to WT (Fig. S6b), smaller than the reduction seen for W99A, which we note as a minor deviation from an otherwise clean biochemical profile. These results identify W99A as an overtly loss-of-function allele, and establish W99F as a biochemically clean, minimally perturbing substitution suitable for chemical-genetic use.

### Cellular target engagement and genetic epistasis confirm on-target activity of CHI-DEL

To determine whether the structure-guided resistance allele of **CHI-DEL** confers resistance at the cellular level, not just biochemically, we compared CETSA in HEK293T cells expressing wild-type or W99F CHI3L1. Before treatment, the isogenic WT, CHI3L1-KO, KO+WT-rescue, and KO+W99F-rescue lines used in the epistasis experiments below were confirmed to be baseline-matched across CHI3L1 level, basal signaling, and baseline invasion/proliferation (Fig. S7). **CHI-DEL** (1 µM) shifted the melt curve in WT-expressing cells relative to vehicle, but produced no shift in W99F-expressing cells, which overlapped closely with vehicle in both genotypes (Fig. 3a). Isothermal dose-response CETSA confirmed this was concentration-dependent: **CHI-DEL** engaged WT CHI3L1 with EC_50_ = 0.24 µM, while W99F showed no concentration-dependent engagement across the same range (Fig. 3b), consistent with the Trp99-dependent binding mode defined structurally in Figs. 1 and 2. Consistent with this cellular resistance, SPR analysis showed no concentration-dependent binding of **CHI-DEL** to purified W99F CHI3L1 across concentrations up to 5 µM (Fig. S8).

**Figure 3.**
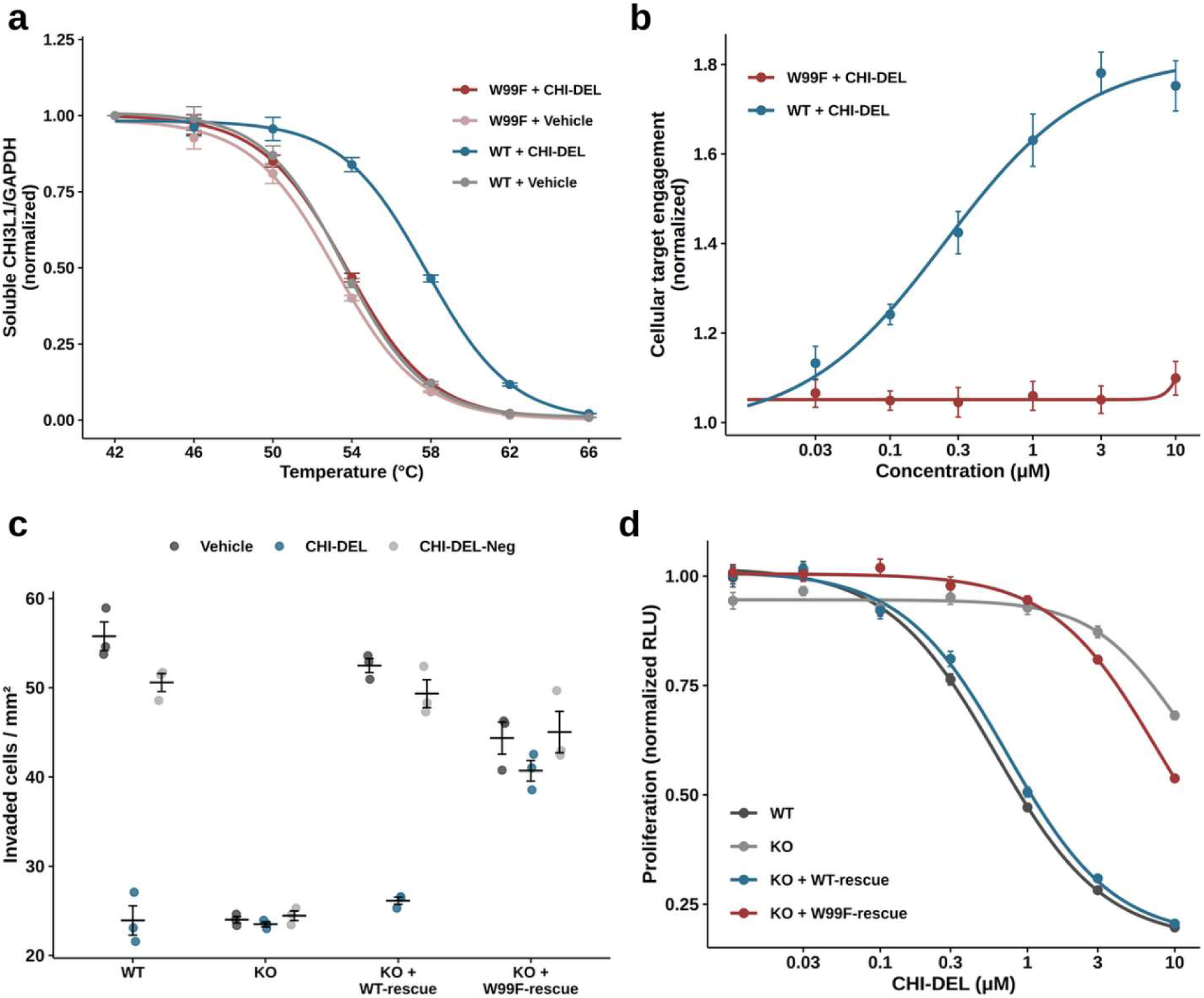
Cellular target engagement and genetic epistasis confirm on-target activity of CHI-DEL. **(a)** CETSA temperature-shift assay in HEK293T cells expressing WT or W99F CHI3L1. **CHI-DEL** (1 µM) shifts the melt curve in WT-expressing cells relative to vehicle, but produces no shift in W99F-expressing cells. **(b)** Isothermal dose-response CETSA; **CHI-DEL** engages WT CHI3L1 with EC_50_ = 0.24 µM but shows no concentration-dependent engagement of W99F. **(c)** Invasion assay across an isogenic HEK293T genotype panel (WT, CHI3L1-KO, KO+WT-rescue, KO+W99F-rescue) and three treatments (Vehicle, **CHI-DEL**, **CHI-DEL-Neg**). **CHI-DEL** suppresses invasion in WT and KO+WT-rescue but not in KO or KO+W99F-rescue, while both rescue genotypes restore baseline (Vehicle) invasion equivalently, confirming W99F retains normal CHI3L1 function. **(d)** Proliferation dose-response to **CHI-DEL** across the same four genotypes; KO+WT-rescue closely tracks WT sensitivity, while KO and KO+W99F-rescue are both right-shifted by approximately an order of magnitude. Data are mean ± SEM from three independent biological experiments (n = 3).

We next tested whether this resistance held in a genetic loss-of-function/rescue system, using an isogenic panel of CHI3L1-knockout HEK293T cells reconstituted with either WT or W99F CHI3L1. In an invasion assay, CHI3L1-KO cells showed reduced invasion relative to WT; reconstitution with WT CHI3L1 restored invasion to WT levels, as did reconstitution with W99F, confirming that W99F retains normal CHI3L1 function in this cellular context (Fig. 3c). **CHI-DEL** suppressed invasion in WT and KO+WT-rescue cells, but had no effect in CHI3L1-KO cells or KO+W99F-rescue cells, while the inactive analog **CHI-DEL-Neg** had no effect in any genotype (Fig. 3c). This same epistasis pattern was reproduced in an independent functional assay: proliferation dose-response to **CHI-DEL** closely tracked between WT and KO+WT-rescue cells, while KO and KO+W99F-rescue cells were both right-shifted by approximately an order of magnitude (Fig. 3d). These results establish that the cellular and functional effects of **CHI-DEL** require the Trp99-dependent binding mode defined structurally in Figure 2, rather than an off-target consequence of compound treatment: a genetic manipulation (CHI3L1 knockout) and a single point mutation (W99F) that leaves native CHI3L1 function intact both abolish **CHI-DEL**’s functional activity, while leaving the underlying WT-like phenotype fully restored.

### CHI-DEL disrupts CHI3L1-CD44 engagement and a multi-node signaling program in a Trp99-dependent manner

To determine whether the structural binding mode of **CHI-DEL** translates into disruption of CHI3L1’s physiological receptor engagement, we used a partner-competition ELISA comparing WT and W99F CHI3L1 in the presence of **CHI-DEL** or **CHI-DEL-Neg**. **CHI-DEL** potently disrupted the CHI3L1-CD44 complex in WT protein (IC_50_ = 0.090 ± 0.009 µM), whereas its effect on W99F was right-shifted and more closely resembled the two **CHI-DEL-Neg** conditions, consistent with loss of engagement at the mutated residue (Fig. 4a). All four conditions showed a nonspecific decline at the highest tested concentration (30 µM), including both **CHI-DEL-Neg** controls. We next asked whether this receptor-engagement effect extended to downstream signaling, and specifically whether it was restricted to pSTAT3, the node most commonly reported for CHI3L1 in the literature. In WT cells, **CHI-DEL** produced a dose-dependent reduction in both pSTAT3/STAT3 and phospho-p65/total-p65 (an NF-κB pathway readout), with near-complete suppression of both nodes at 3 µM; **CHI-DEL-Neg** (3 µM) had no effect on either node (Fig. 4b). This establishes that the signaling effect of **CHI-DEL** is not confined to STAT3 but extends to a second, independent pathway node.

**Figure 4.**
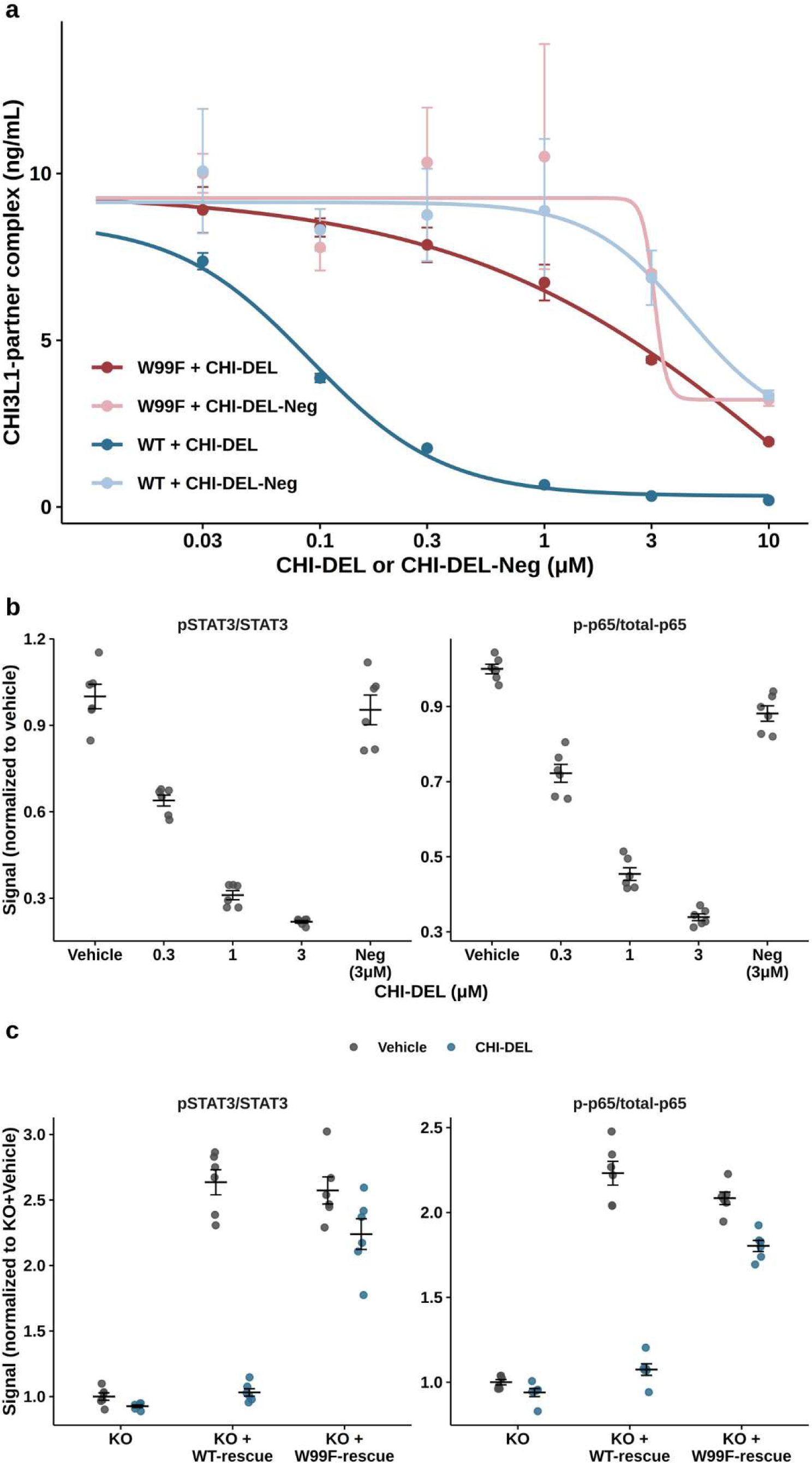
CHI-DEL disrupts CHI3L1-CD44 engagement and multi-node signaling in a Trp99-dependent manner. **(a)** Partner-competition ELISA measuring the CHI3L1-CD44 complex in the presence of **CHI-DEL** or **CHI-DEL-Neg**, for WT and W99F CHI3L1. **CHI-DEL** potently disrupts the complex in WT protein; its effect on W99F is right-shifted and resembles the **CHI-DEL-Neg** conditions. **(b)** Dose-dependent effect of **CHI-DEL** on two signaling nodes, pSTAT3/STAT3 and phospho-p65/total-p65, in WT CHI3L1-expressing cells. **CHI-DEL-Neg** (3 µM) has no effect on either node. **(c)** The same two signaling nodes in an isogenic CHI3L1-knockout/rescue panel (KO, KO+WT-rescue, KO+W99F-rescue), each with Vehicle or **CHI-DEL** (1 µM). **CHI-DEL** collapses both nodes to near-baseline in KO+WT-rescue cells, but its effect in KO+W99F-rescue cells is attenuated rather than abolished. Data are mean ± SEM from three independent biological experiments (n = 3).

To test whether this multi-node signaling effect requires the Trp99-dependent binding mode, we examined the same two nodes in the isogenic CHI3L1-knockout/rescue panel. **CHI-DEL** collapsed both pSTAT3 and p-p65 to near-baseline levels in KO+WT-rescue cells, closely matching the untreated KO condition. In KO+W99F-rescue cells, however, **CHI-DEL**’s effect on both nodes was substantially attenuated rather than abolished, with signal remaining well above the KO baseline (Fig. 4c). This partial, rather than complete, resistance at the signaling level is a more modest effect than the essentially complete resistance observed functionally in Figure 3, and is reported here without overstating its magnitude. These data indicate that **CHI-DEL** disrupts CHI3L1’s physiological receptor engagement and a multi-node downstream signaling program, extending beyond the commonly reported pSTAT3 axis, in a manner that depends substantially, though not completely, on the Trp99-defined binding site established structurally in Figure 2.

### CHI3L1-genotype-dependent GBM signaling drives a reversible microglial phenotype

To determine whether the mechanism of **CHI-DEL** extends to a disease-relevant, non-transformed cellular context, we exposed human iPSC-derived microglia to conditioned medium from an isogenic panel of CHI3L1-genotyped GBM cells (WT, CHI3L1-knockout, KO+WT-rescue, and KO+W99F-rescue), combined with direct compound treatment of the microglia, yielding a 9-arm design that mirrors the genetic epistasis logic of Figs. 3 and 4. Conditioned medium from WT CHI3L1-expressing GBM cells induced a polarized phenotype in the recipient microglia relative to medium from control (non-GBM) cells, evident functionally (increased feedback invasion of GBM cells, *t* = 10.8, *p* = 0.0012, *n* = 4), by surface marker (CD206⁺ frequency, *t* = 25.3, *p* < 0.0001, *n* = 3 independent differentiations), by gene expression (SPP1 mRNA, *t* = 13.8, *p* = 0.0047, *n* = 3; ARG1 mRNA, *t* = 17.3, *p* = 0.0021, *n* = 3, Fig. S9), and by secreted cytokine (IL-10, *t* = 19.1, *p* < 0.0001, *n* = 4) (Fig. 5a–d). Direct **CHI-DEL** treatment of the microglia reversed this phenotype across all five readouts, while **CHI-DEL-Neg** did not (all comparisons *p* < 0.01, including ARG1, *t* = 53.8, *p* < 0.0001, Fig. S9), and conditioned medium from CHI3L1-knockout GBM cells alone phenocopied the drug-treated state.

**Figure 5.**
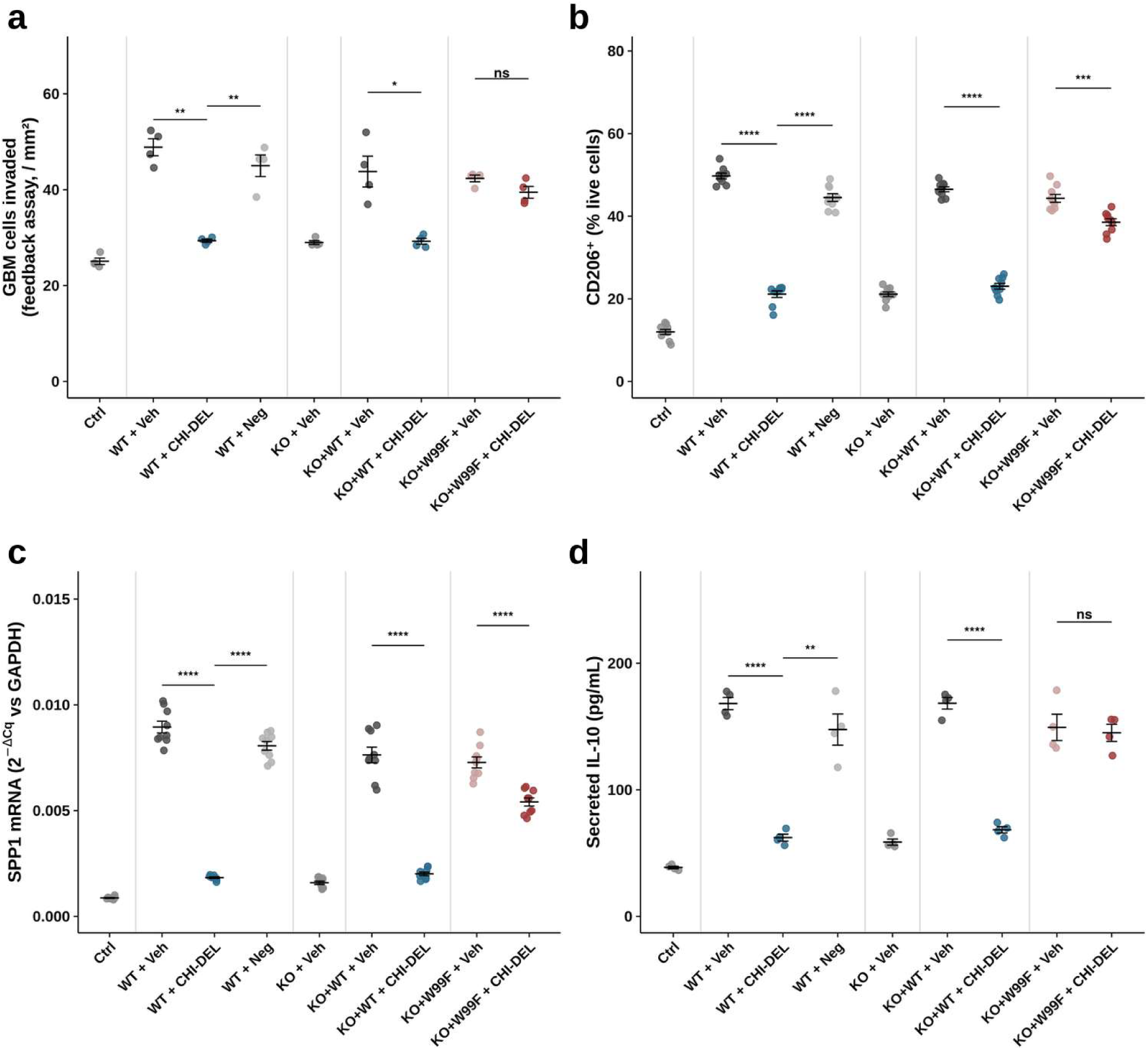
CHI3L1-genotype-dependent GBM conditioned medium drives a reversible, Trp99-dependent phenotype in human iPSC-derived microglia. **(a)** Functional feedback invasion assay: GBM cell invasion in response to conditioned medium from iPSC-derived microglia, across microglia exposed to conditioned medium from a 9-arm panel of CHI3L1-genotyped GBM cells (WT, CHI3L1-KO, KO+WT-rescue, KO+W99F-rescue) with direct microglial compound treatment (n = 4 biological replicates/arm). (**b)** CD206⁺ frequency by flow cytometry (% live cells) in the recipient microglia; small points show individual replicate wells, triangles show the mean of each independent differentiation batch (*n* = 3 batches/arm), the unit used for statistical testing. **(c)** SPP1 mRNA by qPCR in the recipient microglia, shown with the same well-level/run-level convention as **(b)** (n = 3 independent runs/arm); ARG1 mRNA from the same runs is shown in Fig. S9. **(d)** Secreted IL-10 by ELISA from the recipient microglia (n = 4 biological replicates/arm). In all panels, thin vertical lines demarcate genotype blocks (Control | WT-conditioned medium | CHI3L1-KO-conditioned medium | KO+WT-rescue | KO+W99F-rescue); data are mean ± SEM. Brackets show two-sided Welch’s *t*-tests for four mechanism-defining comparisons per panel; ns, not significant; \**p* < 0.05, \*\**p* < 0.01, \*\*\**p* < 0.001, \*\*\*\**p* < 0.0001; exact *p*-values, *t*-statistics, and degrees of freedom for all comparisons (including Fig. S9) are reported in the source data.

Reconstituting the knockout GBM cells with WT CHI3L1 restored both the polarized phenotype and **CHI-DEL** sensitivity in the recipient microglia across all five readouts (all *p* < 0.003, including ARG1, Fig. S9). Reconstitution with W99F CHI3L1 restored the baseline polarized phenotype equivalently to WT-rescue, but the effect of **CHI-DEL** on microglia exposed to this conditioned medium was consistently smaller and graded across readouts. Functional invasion (*t* = 2.05, *p* = 0.098, *n* = 4) and secreted IL-10 (*t* = 0.35, *p* = 0.74, *n* = 4) showed no statistically significant **CHI-DEL** response with W99F-rescue-conditioned medium, indicating essentially complete resistance at these two levels. In contrast, CD206 (*t* = 3.02, *p* = 0.041, *n* = 3), SPP1 (*t* = 13.9, *p* < 0.001, *n* = 3), and ARG1 (*t* = 5.17, *p* = 0.035, *n* = 3, Fig. S9) each showed a statistically significant but visibly attenuated reduction relative to the corresponding WT-rescue comparison. Across all five readouts, then, the W99F mutation in the GBM-cell source does not uniformly abolish the downstream effect of **CHI-DEL** on microglia, but substantially blunts it, completely at the functional and secreted-cytokine level, partially at the surface-marker and transcript level, consistent with the partial signaling resistance already observed in Figure 4.

### In vitro ADME and permeability profiling of CHI-DEL

To assess whether the physicochemical and metabolic properties of **CHI-DEL** were compatible with systemic and CNS exposure, we evaluated its lipophilicity, solubility, permeability, metabolic stability and plasma protein binding (Table S2). **CHI-DEL** exhibited a LogD_7.4_ of 2.63 and kinetic solubility of 128 µM in PBS containing 1% DMSO. In PAMPA, **CHI-DEL** showed a P_app_ of 5.2 × 10^-6^ cm/s, while bidirectional MDCK-MDR1 assays yielded P_app_ values of 7.9 × 10^-6^ cm/s in the A→B direction and 10.1 × 10^-6^ cm/s in the B→A direction, corresponding to an efflux ratio of 1.28. Addition of elacridar reduced the efflux ratio to 1.10, consistent with limited P-gp-mediated efflux. **CHI-DEL** was metabolically stable in mouse and human liver microsomes, with half-lives of 96 ± 8 and 121 ± 11 min, respectively, and remained stable in both mouse and human plasma for more than 180 min. Human plasma protein binding was 92.8 ± 0.7%. **CHI-DEL** also showed minimal inhibition of major CYP isoforms and weak hERG liability. Collectively, these properties are consistent with a profile compatible with systemic exposure and CNS penetration.

### CHI-DEL is brain- and tumor-penetrant and produces dose-dependent, on-target pharmacodynamic effects in vivo

To determine whether the target engagement and mechanism of **CHI-DEL** translate to an intact, brain tumor-bearing animal, we first characterized its pharmacokinetics in tumor-bearing mice. Following a single 20 mg/kg dose, **CHI-DEL** was rapidly absorbed and distributed to both brain and tumor tissue at concentrations tracking closely with plasma across the full 24-hour profile, confirming brain penetrance sufficient to support intracranial target engagement (Fig. 6a). Brain:plasma and tumor:plasma ratios increased over the 24 h profile, consistent with tissue retention (Fig. S10).

**Figure 6.**
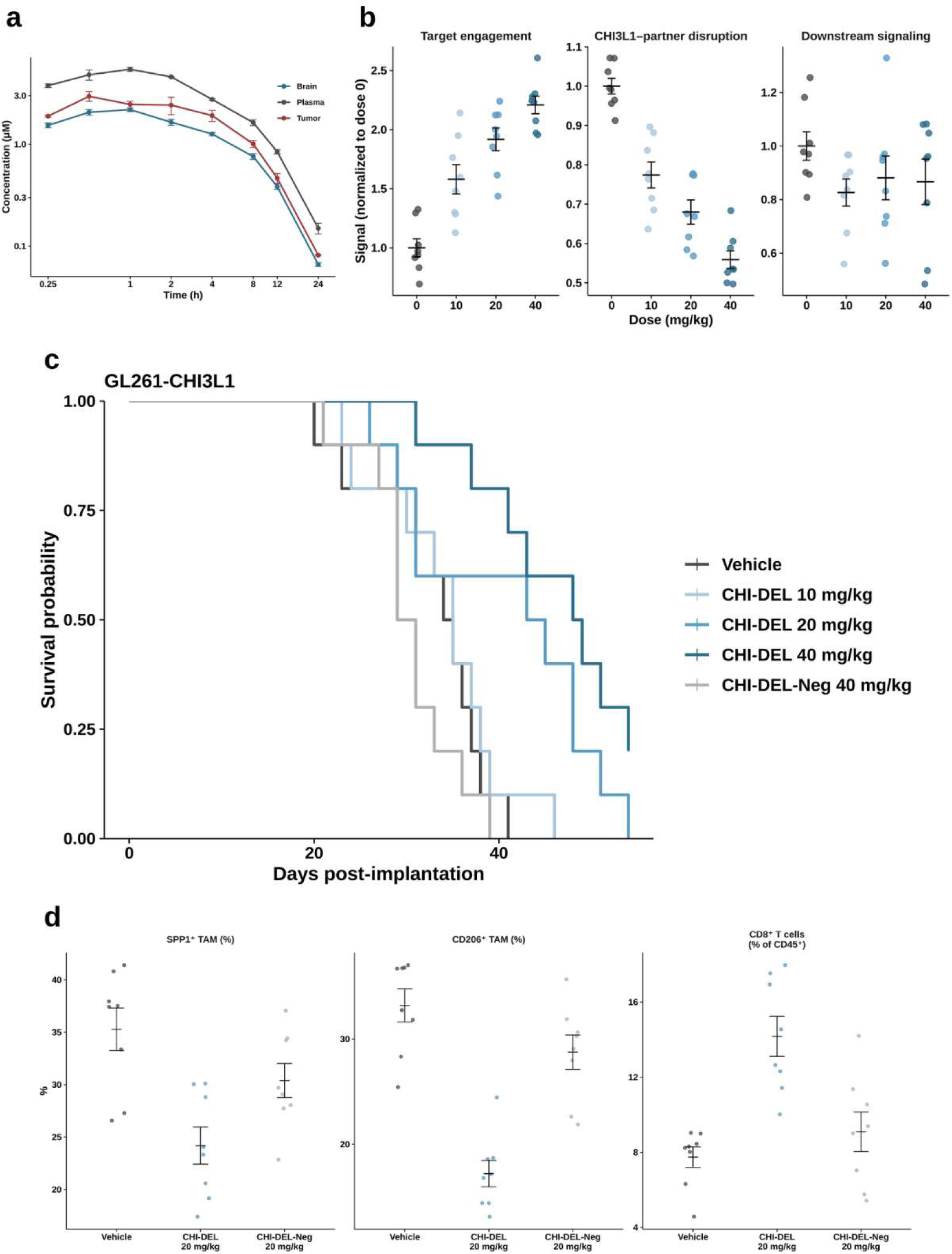
CHI-DEL is brain-penetrant, engages CHI3L1 in tumor tissue in a dose-dependent manner, and extends survival with immune repolarization. **(a)** Pharmacokinetic profile of **CHI-DEL** (20 mg/kg, single dose) in plasma, brain, and tumor tissue over 24 h (n = 3 mice per time point). Data are mean ± SEM. **(b)** Dose-dependent tumor pharmacodynamics in mice bearing orthotopic GL261-CHI3L1 tumors (n = 8/dose): target engagement (CETSA soluble CHI3L1 fraction, normalized to dose 0), CHI3L1-CD44 disruption (partner-complex ELISA, normalized to dose 0), and downstream signaling (pSTAT3/STAT3, normalized to dose 0). Tumor tissue was collected 4 h after the final scheduled dose. **(c)** Kaplan-Meier survival in the GL261-CHI3L1 orthotopic model (n = 10/group; 5F/5M): vehicle, **CHI-DEL** (10, 20, or 40 mg/kg), and **CHI-DEL-Neg** (40 mg/kg). Log-rank p = 3.6 × 10^-4^ (vehicle vs. **CHI-DEL** 40 mg/kg); p = 5.5 × 10^-5^ (**CHI-DEL** 40 mg/kg vs. **CHI-DEL-Neg** 40 mg/kg). **(d)** Tumor immune profiling by flow cytometry in GL261-CHI3L1 tumors (n = 8/group): SPP1⁺ and CD206⁺ frequency among tumor-associated macrophages, and CD8⁺ T cell frequency among CD45⁺ cells. **CHI-DEL** reduces both immunosuppressive TAM markers and increases CD8⁺ T cell infiltration relative to vehicle; **CHI-DEL-Neg** has no effect on any of the three readouts. Tumor tissue was collected 4 h after the final scheduled dose. Data are mean ± SEM in all panels; each point is one animal.

We then assessed dose-dependent tumor target engagement in mice bearing orthotopic GL261-CHI3L1 tumors, dosed twice weekly with **CHI-DEL** (10, 20, or 40 mg/kg), with tumors collected 4 h after the final scheduled dose. **CHI-DEL** produced a clear, dose-dependent increase in the soluble (non-aggregated) fraction of tumor CHI3L1 by CETSA, consistent with direct target engagement in tumor tissue, and a corresponding dose-dependent disruption of the CHI3L1-CD44 complex (Fig. 6b). Downstream pSTAT3/STAT3 signaling showed a modest reduction at the lowest dose but did not decline further with increasing dose and displayed substantial variability at every dose tested (Fig. 6b), in contrast to the clean dose-response seen for the two more proximal engagement readouts. This dissociation, robust, dose-dependent target engagement without a correspondingly clean downstream signaling readout in bulk tumor tissue, may reflect the cellular heterogeneity of tumor tissue relative to the pure cell populations used in Figs 1-4, and is discussed further below.

### CHI-DEL extends survival in an orthotopic glioma model

To test whether this target engagement translated to therapeutic benefit, we assessed survival in mice bearing orthotopic GL261-CHI3L1 tumors treated with **CHI-DEL** across the same three doses, alongside vehicle and an inactive-analog control (**CHI-DEL-Neg**, 40 mg/kg). **CHI-DEL** extended survival in a dose-dependent manner, with the highest dose (40 mg/kg) producing the greatest benefit (log-rank vs. vehicle, *p* = 3.6 × 10^-4^; *n* = 10/group), while **CHI-DEL-Neg** at the same 40 mg/kg dose provided no benefit over vehicle and was significantly less effective than CHI-DEL itself (log-rank, *p* = 5.5 × 10^-5^), confirming that this benefit depends on the specific compound rather than a generic effect of dosing (Fig. 6c). **CHI-DEL** was well tolerated at all doses tested, with no meaningful body-weight loss over 21 days of repeat dosing (Fig. S11).

### CHI-DEL remodels the tumor immune microenvironment in vivo

Given the established role of CHI3L1 in shaping an immunosuppressive tumor microenvironment, we profiled tumor-associated macrophages (TAMs) and T cells by flow cytometry in GL261-CHI3L1 tumors from mice treated with vehicle, **CHI-DEL** (20 mg/kg), or **CHI-DEL-Neg** (20 mg/kg). **CHI-DEL** treatment reduced both SPP1⁺ and CD206⁺ TAM frequencies relative to vehicle, markers associated with an immunosuppressive, pro-tumorigenic macrophage state, while **CHI-DEL-Neg** produced no change in either marker (Fig. 6d). Concurrently, **CHI-DEL** increased the frequency of CD8⁺ T cells among CD45⁺ immune cells, consistent with a shift toward a less immunosuppressive, more anti-tumor immune microenvironment accompanying CHI3L1 target engagement (Fig. 6d).

## Discussion

This work establishes **CHI-DEL** as a validated, structurally characterized small molecule CHI3L1 ligand and uses a structure-guided resistance mutation to provide direct chemical-genetic proof that its cellular and functional effects are mediated through the specific binding site defined crystallographically. Three aspects of this work are worth discussing on their own terms. First, the dissociation between DNA-encoded library enrichment and off-DNA binding (Fig. 1) demonstrates quantitatively that enrichment magnitude does not predict target engagement: two candidates with 1.2- to 3-fold higher enrichment than **CHI-DEL** both failed to show reproducible SPR binding. This underscores that off-DNA biophysical validation is a necessary filter in DEL campaigns.

Second, the Trp99-centered structural rationale (Fig. 2) proved to be a genuinely predictive design principle rather than a post-hoc explanation. The W99F mutation was designed conservatively to preserve native CHI3L1 biology while disrupting **CHI-DEL** engagement, and it did so: indistinguishable from WT on secretion, basal signaling, and thermal stability, in contrast to the destabilizing W99A control. This qualification step, performed before any functional use of the mutant, is what allows the subsequent epistasis experiments (Figs. 3-5) to be interpreted as evidence of on-target pharmacology rather than an artifact of a partially loss-of-function allele.

Third, W99F produced strong resistance at the level of **CHI-DEL** target engagement, while the magnitude of resistance across downstream molecular and functional readouts varied. Purified W99F CHI3L1 showed no concentration-dependent binding to **CHI-DEL** by SPR across the tested concentration range, and W99F similarly abolished concentration-dependent cellular target engagement by CETSA. Downstream phenotypic resistance was more graded. In the human iPSC-microglia epistasis panel (Fig. 5), resistance was essentially complete for functional invasion and secreted IL-10, whereas CD206, SPP1 and ARG1 retained partial responses to **CHI-DEL**. Partner-complex disruption and downstream signaling (Fig. 4) likewise showed attenuated rather than complete resistance. Thus, disruption of the Trp99-defined binding mode strongly impairs **CHI-DEL** target engagement, while the extent to which this propagates through downstream CHI3L1-dependent signaling and cellular phenotypes varies by biological readout.

The in vivo data extend this work in a different direction than the cellular chemical-genetic experiments. Because the W99F genetic-resistance design was not carried into the animal studies, in vivo validation instead relies on pharmacological specificity, a dose-response relationship and a structurally related but inactive analog, **CHI-DEL-Neg**. This is a legitimate but distinct form of evidence from the genetic proof established in vitro, and we are explicit about that distinction rather than implying the in vivo survival data constitute genetic resistance evidence. Within the GL261-CHI3L1 model, **CHI-DEL** produced dose-dependent tumor target engagement (CETSA, partner-complex disruption) and repolarized the tumor immune microenvironment (reduced SPP1⁺/CD206⁺ TAMs, increased CD8⁺ T cell infiltration), consistent with the neuroimmune mechanism proposed throughout this work. Notably, the downstream pSTAT3 signaling readout, the pathway most directly implicated in vitro (Figs. 1, 4), did not show a clean dose-response in bulk tumor tissue, unlike the more proximal engagement readouts. The absence of a clear dose-response in bulk tumor pSTAT3 may arise from cellular heterogeneity within the tumor, differences between cultured cells and intact tumor tissue, or temporal dissociation between proximal target engagement and downstream signaling.

In summary, our work provides a template for establishing genetic proof of on-target pharmacology for a non-enzymatic, secreted protein target lacking a catalytic site to mutate, using a chemical-genetic resistance allele validated at the structural, biochemical, and cellular level, while separately demonstrating that the resulting compound achieves brain penetrance, dose-dependent tumor target engagement, and survival benefit in an orthotopic glioma model.

## Materials and Methods

### DNA-encoded library screening

A 1.4 billion DNA-Encoded Chemical Library Kit (AlphaMa SDEL Pubkit) was purchased from AlphaMa Biotechnology, Suzhou. One kit afforded one screening condition against either the target or blank control bead as a no-target screening (NTC). DEL screening against CHI3L1 was performed by immobilizing 10 µg of target protein on 20 µL of HisPur™ Ni-NTA magnetic beads and incubating the immobilized protein with the DNA-encoded chemical library. Three rounds of affinity screening were performed in each condition, including protein immobilization, DEL panning, wash steps, and heated elution, following the manufacturer’s instructions. Heated elution samples from the third round of affinity screening were subjected to Next-Generation Sequencing to reveal the identity of the bound compounds under each screening condition. Utilizing AlphaMa’s decoding program (not disclosed), the FASTQ files were converted into chemical identities and the corresponding copies in each screening condition.

Enriched library members and associated building-block combinations were identified by comparing their representation under the CHI3L1 selection and matched no-target control conditions. Compounds displaying CHI3L1-associated enrichment and distinguishable structure-enrichment relationships were prioritized for off-DNA synthesis and subsequent biophysical validation. The chemical characterization and purity assessment of the synthesized compounds are presented in Figs. S12-S21.

### Surface plasmon resonance

Surface plasmon resonance measurements were performed using a Biacore 8K instrument (Cytiva) equipped with a CM5 sensor chip. Recombinant His-tagged human CHI3L1 was diluted to 8 µg/ml in immobilization buffer at pH 5.0 and covalently immobilized on the active flow cells by amine coupling to approximately 3,300-3,500 response units (RU). Reference flow cells underwent the same activation and blocking procedure without protein immobilization.

Off-DNA compounds were initially evaluated using single-cycle kinetics with PBS-P containing 2.5% (v/v) DMSO as the running buffer. Eight sequential concentrations of each compound (0.091, 0.27, 0.82, 2.47, 7.41, 22.22, 66.67 and 200 µM) were injected in ascending order without regeneration between injections. Each concentration was injected for 120 s at a flow rate of 30 µl/min, followed by a final dissociation phase of 600 s. Buffer-only injections and solvent-correction cycles were included. The sensor surface was regenerated between compound cycles using 10 mM sodium acetate containing 250 mM NaCl, pH 4.0, for 30 s at a flow rate of 30 µl/min. Selected compounds were subsequently reassessed in independent experiments using compound-specific concentration ranges. **CHI-DEL** was tested at 0.0091, 0.027, 0.082, 0.25, 0.74, 2.22, 6.67 and 20 µM.

**CHI-DEL** selectivity (at a single concentration of 50 µM) was assessed by SPR against a panel of 15 related and unrelated recombinant proteins, including the chitinase/chitinase-like proteins CHI3L2, CHIT1 and CHIA and CHI3L1-associated proteins RAGE, galectin-3, IL13Rα2 and CD44. Proteins were immobilized on CM5 sensor chips. Because the counterscreen was intended to identify appreciable off-target binding rather than derive affinity values for nonbinding proteins, quantitative K_D_ values were not assigned to surfaces that failed to produce concentration-dependent, saturable binding. Responses are reported as reference-corrected RU. Sensorgrams were corrected using the corresponding reference-surface responses, buffer-only injections and solvent-correction data and analyzed using Biacore Insight Evaluation software (version 6.0.7.1750). Corrected sensorgrams were fitted to a 1:1 binding model to estimate the association-rate constant (k_a_) and dissociation-rate constant (k_d_). The equilibrium dissociation constant was calculated as K_D_ = k_d_/k_a_. Compounds were advanced only when they generated concentration-dependent sensorgrams that were reproducible across independent experiments and adequately described by the binding model. Affinity estimates derived from low-response, irregular, poorly fitted or non-reproducible profiles were not used for compound prioritization.

### Expression and purification of recombinant human CHI3L1

Human CHI3L1 (residues 1-383) was cloned into a pCMV expression vector with a C-terminal HRV 3C protease cleavage site followed by a polyhistidine tag. Recombinant CHI3L1 was transiently expressed in Expi293F™ suspension cells using the Expi293™ Expression System (Thermo Fisher Scientific). Cells were maintained in Expi293™ Expression Medium at 37 °C under 8% CO_2_ with orbital shaking according to the manufacturer’s recommendations. Transfection was performed using the ExpiFectamine™ 293 Transfection Kit at a plasmid DNA-to-transfection reagent ratio of 1:5. Culture supernatants were harvested 5 days after transfection.

The supernatants were clarified by centrifugation and incubated with manually packed Ni-NTA resin pre-equilibrated in binding buffer containing 20 mM Tris-HCl, pH 8.0, 150 mM NaCl, 5% (v/v) glycerol and 10 mM imidazole. After extensive washing with binding buffer, bound CHI3L1 was eluted using the same buffer supplemented with 300 mM imidazole. Fractions containing CHI3L1 were pooled and further purified by size-exclusion chromatography in 20 mM Tris-HCl, pH 8.0, 150 mM NaCl and 5% (v/v) glycerol. Peak fractions were pooled and concentrated to approximately 5 mg ml^-1^ for crystallization.

### Crystallization, X-ray data collection and structure determination

Purified recombinant human CHI3L1 was concentrated to 5 mg ml^-1^ in 20 mM Tris-HCl, pH 8.0, 150 mM NaCl and 5% (v/v) glycerol. CHI3L1 was mixed with **CHI-DEL** at a protein-to-ligand molar ratio of 1:3 before crystallization. Crystals of the CHI3L1-**CHI-DEL** complex were grown at 289 K using the sitting-drop vapor-diffusion method by mixing the protein-ligand solution with reservoir solution containing 0.1 M HEPES, pH 7.0, and 10% (w/v) PEG 6000. Initial crystal hits were observed over a pH range of 5.0-7.0 and subsequently optimized to obtain crystals suitable for diffraction analysis.

X-ray diffraction data were collected at 100 K on beamline 17-ID-1 at the National Synchrotron Light Source II (NSLS-II) using a Dectris EIGER X 9M detector and an X-ray wavelength of 0.97934 Å. Diffraction images were indexed, integrated and scaled using autoPROC and AIMLESS ^21,22^. The structure was determined by molecular replacement with PHASER using the previously reported CHI3L1 structure (PDB 8R41) as the search model ^23^. Iterative model building and refinement were performed using Coot and PHENIX version 1.21.1 ^24,25^. Data-collection and refinement statistics are provided in Table S1.

### Structure-guided mutagenesis

Candidate resistance substitutions were selected from residues contacting or immediately surrounding **CHI-DEL** in the 2.81-Å CHI3L1-**CHI-DEL** co-crystal structure (PDB 10JV). Mutations were introduced into the human CHI3L1 expression construct using the Q5 Site-Directed Mutagenesis Kit (New England Biolabs) according to the manufacturer’s protocol. Mutagenic primers were designed to introduce the desired codon substitution without altering neighboring residues. PCR products were kinase-ligase-DpnI treated, transformed into chemically competent *Escherichia coli*, and individual colonies were expanded for plasmid purification. The complete CHI3L1 coding sequence of each construct was verified by Sanger sequencing before experimental use. WT and mutant constructs were propagated and prepared using the same plasmid purification procedure. WT, W99F, W99A, Y206F and R263A CHI3L1 were evaluated before their use in chemical-genetic experiments to distinguish selective resistance to **CHI-DEL** from nonspecific impairment of protein expression, secretion, stability or biological activity.

### CHI3L1 secretion

HEK293T cells were maintained in DMEM supplemented with 10% fetal bovine serum and 1% penicillin-streptomycin at 37 °C in a humidified atmosphere containing 5% CO_2_. Cells were seeded 24 h before transfection and transfected at approximately 70-80% confluence with equal amounts of WT, W99F, W99A, Y206F or R263A CHI3L1 plasmid using Lipofectamine 3000. Six hours after transfection, the medium was replaced with fresh complete medium. After an additional 24 h, cells were washed twice with PBS and transferred to serum-reduced medium containing 1% fetal bovine serum. Conditioned medium was collected 24 h later, corresponding to 48 h after transfection, and clarified by centrifugation at 1,000 × g for 10 min at 4 °C.

Secreted CHI3L1 was quantified using Human Chitinase 3-like 1/YKL-40 Quantikine ELISA Kit (R&D Systems, Cat. No. DC3L10) according to the manufacturer’s instructions. Conditioned-medium samples were diluted within the validated dynamic range of the assay and measured in duplicate. CHI3L1 concentrations were normalized to total cellular protein determined from the corresponding producer-cell lysates using a BCA assay. Nine independent biological replicates were analyzed per genotype.

### Basal CHI3L1-dependent signaling

To determine whether the candidate resistance mutations preserved basal CHI3L1-dependent signaling, HEK293T cells expressing WT or mutant CHI3L1 were harvested 48 h after transfection. Cells were washed twice with ice-cold PBS and lysed in buffer supplemented with protease and phosphatase inhibitor cocktails. Lysates were clarified at 16,000 × *g* for 15 min at 4 °C, and total protein concentration was determined by BCA assay. Phospho-STAT3 (Tyr705) and total STAT3 were quantified using the STAT3 (pY705) + total STAT3 ELISA Kit (Abcam, ab126459) according to the manufacturer’s instructions. Equivalent amounts of total protein were analyzed for each sample. Phospho-STAT3 signal was normalized to total STAT3 and expressed relative to the CHI3L1-knockout baseline. Nine independent biological replicates were analyzed per genotype.

### Differential scanning fluorimetry

Recombinant WT, W99F, W99A, Y206F and R263A CHI3L1 proteins were diluted to 5 µM in 20 mM Tris-HCl, pH 8.0, 150 mM NaCl and 5% glycerol. Protein was mixed with SYPRO Orange dye at a final 5× concentration in 20-µl reactions in optical 96-well plates. Fluorescence was recorded on a real-time PCR instrument while the temperature was increased from 25 to 95 °C at approximately 1 °C min⁻¹. Apparent melting temperatures were calculated from the maximum of the first derivative of the fluorescence-versus-temperature curve. Nine independent protein preparations were analyzed per variant. Each preparation was measured in technical triplicate, and the technical measurements were averaged to yield one value per biological replicate (n = 9 per variant).

W99F, Y206F and R263A were considered structurally tolerated because their apparent melting temperatures differed from WT by no more than approximately 1 °C, whereas W99A exhibited an approximately 7 °C decrease in apparent thermal stability together with reduced secretion and basal signaling. The latter was therefore classified as a destabilizing loss-of-function control rather than a suitable resistance allele.

### Cellular thermal shift assay

Cellular target engagement of **CHI-DEL** was evaluated using temperature-response and isothermal dose-response CETSA. HEK293T cells expressing WT or W99F CHI3L1 were treated with vehicle, 1 µM **CHI-DEL** or 1 µM **CHI-DEL-Neg** for 1 h at 37 °C. Cells were washed twice with PBS, detached using enzyme-free cell dissociation buffer, collected by centrifugation and resuspended in PBS supplemented with protease inhibitors. Equal cell suspensions were divided into PCR tubes and heated for 3 min at temperatures ranging from 40 to 64 °C in 3 °C increments, followed by cooling for 3 min at room temperature. Cells were lysed by three freeze-thaw cycles consisting of freezing in liquid nitrogen and thawing at 25 °C. Lysates were centrifuged at 20,000 × g for 20 min at 4 °C to pellet thermally aggregated protein. Soluble CHI3L1 in the post-centrifugation supernatant was quantified using a Human Chitinase 3-like 1/YKL-40 Quantikine ELISA Kit (R&D Systems, Cat. No. DC3L10) according to the manufacturer’s instructions. CHI3L1 levels were normalized to the lowest-temperature sample within each genotype and treatment condition. Thermal-denaturation curves were fitted by nonlinear regression using a variable-slope sigmoidal model, and apparent melting temperatures were derived from the fitted curves.

For isothermal dose-response CETSA, WT- or W99F-expressing HEK293T cells were incubated with **CHI-DEL** or **CHI-DEL-Neg** over a concentration range of 0.01-10 µM for 1 h at 37 °C. Cells were then heated at 55 °C for 3 min, cooled for 3 min, lysed by three freeze-thaw cycles and centrifuged at 20,000 × g for 20 min at 4 °C. Soluble CHI3L1 was quantified as mentioned above. Concentration-dependent stabilization was analyzed using four-parameter logistic nonlinear regression. EC_50_ values were calculated only when the data supported a monotonic concentration-dependent response. Three independent biological experiments were performed.

### Generation of CHI3L1-knockout and rescue cell systems

CHI3L1-knockout cells were generated using CRISPR-Cas9 editing. For the cellular chemical-genetic experiments in Figure 3, an isogenic HEK293T system was used. A single-guide RNA targeting an early coding exon of human CHI3L1 was cloned into a Cas9-expressing plasmid containing a puromycin-resistance cassette. Cells were transfected with the CRISPR construct and selected with puromycin for 48 h beginning 24 h after transfection. Single cells were subsequently isolated by limiting dilution. Candidate knockout clones were screened by PCR amplification of the targeted region followed by Sanger sequencing and were further validated by loss of CHI3L1 protein in both cellular lysates and conditioned medium.

For rescue experiments, CHI3L1-knockout cells were transduced with lentiviral vectors encoding WT CHI3L1 or W99F CHI3L1 under the same promoter. Stable populations were selected with blasticidin and subsequently maintained without selection antibiotic during functional experiments. WT- and W99F-rescue populations were selected to provide comparable CHI3L1 expression and secretion, as determined by ELISA. The final isogenic panel used for chemical-genetic analysis consisted of parental WT cells, CHI3L1-knockout cells, CHI3L1-knockout cells rescued with WT CHI3L1 and CHI3L1-knockout cells rescued with W99F CHI3L1.

### Cell invasion assay

Cell invasion was measured using 24-well Transwell inserts with 8-µm pores coated with growth-factor-reduced Matrigel. Cells were serum-starved for 6 h before the assay. A total of 5 × 10^4^ cells were suspended in 200 µl serum-free medium and placed in the upper chamber. The lower chamber contained 600 µl medium supplemented with 10% fetal bovine serum as chemoattractant. Cells were treated with vehicle, 1 µM **CHI-DEL** or 1 µM **CHI-DEL-Neg** at the time of plating and allowed to invade for 20 h at 37 °C. Noninvading cells were removed from the upper surface of the membrane with a cotton swab. Cells that had migrated to the lower surface were fixed with 4% paraformaldehyde for 15 min and stained with 0.1% crystal violet.

Membranes were washed extensively, and invaded cells were quantified by imaging five randomly selected fields per insert using identical acquisition settings. Field-level values were averaged to obtain a single value for each insert. Technical replicate inserts were subsequently averaged to provide one value per independent biological experiment. The chemical-genetic comparisons were WT ± **CHI-DEL**, CHI3L1-KO ± **CHI-DEL**, KO+WT rescue ± **CHI-DEL** and KO+W99F rescue ± **CHI-DEL**. **CHI-DEL-Neg** was included throughout as the pharmacological negative control.

### Cell proliferation assay

Cells were seeded in 96-well plates at 3,000 cells per well and allowed to attach overnight. The following day, cells were treated with a logarithmic concentration series of **CHI-DEL** ranging from 3 nM to 30 µM. DMSO concentration was held constant at 0.1% in all wells. After 72 h, cellular ATP content was quantified using CellTiter-Glo 2.0 according to the manufacturer’s instructions. Luminescence was measured after 10 min of equilibration at room temperature. Responses were normalized to vehicle-treated cells within each genotype to account for differences in untreated growth rate. Concentration-response curves were fitted using a four-parameter logistic model with variable slope, and IC_50_ values with 95% confidence intervals were calculated when the data permitted reliable curve fitting. Three independent experiments were performed, each containing three technical wells per concentration.

### Generation of the GBM CHI3L1-knockout and rescue panel

The GBM knockout/rescue panel used for the conditioned-medium experiments was generated from human U87MG glioblastoma cells obtained from the American Type Culture Collection (ATCC HTB-14). Cells were maintained in DMEM supplemented with 10% fetal bovine serum and 1% penicillin-streptomycin at 37 °C in a humidified atmosphere containing 5% CO_2_. Cell identity was confirmed by short-tandem-repeat profiling, and cultures were routinely confirmed to be free of mycoplasma contamination.

CHI3L1-knockout cells were generated using CRISPR-Cas9 editing with a single-guide RNA targeting an early coding exon of human CHI3L1. Cells were transfected with a Cas9 and single-guide RNA expression plasmid containing a puromycin-resistance cassette and selected with 2 µg ml^-1^ puromycin for 48 h beginning 24 h after transfection. Single-cell clones were isolated by limiting dilution. Candidate knockout clones were verified by PCR amplification and Sanger sequencing of the targeted locus and by loss of CHI3L1 protein in both cell lysates and conditioned medium.

For rescue experiments, a validated CHI3L1-knockout clone was transduced with lentiviral vectors encoding wild-type human CHI3L1 or W99F CHI3L1 under the control of the EF1α promoter. Stable populations were selected with 5 µg ml^-1^ blasticidin and subsequently maintained without selection antibiotic during functional experiments. Wild-type- and W99F-rescue populations were matched for cellular and secreted CHI3L1 abundance by ELISA. The final isogenic panel comprised parental wild-type cells, CHI3L1-knockout cells, knockout cells rescued with wild-type CHI3L1, and knockout cells rescued with W99F CHI3L1.

### CHI3L1-CD44 competition assay

Disruption of CHI3L1 receptor/partner engagement was measured using a plate-based competition ELISA. Recombinant human CD44 extracellular domain was used as the CHI3L1 binding partner. High-binding 96-well plates were coated overnight at 4 °C with recombinant human CD44 at 2 µg ml^-1^ in carbonate-bicarbonate buffer, pH 9.6. Plates were washed with PBS containing 0.05% Tween-20 and blocked for 1 h at room temperature with 3% BSA in PBS. Recombinant WT or W99F CHI3L1 was preincubated for 30 min at room temperature with vehicle, **CHI-DEL** or **CHI-DEL-Neg** over a concentration range of 0.01-30 µM in PBS containing 0.1% BSA. Protein-compound mixtures were subsequently transferred to CD44-coated wells and incubated for 1 h at room temperature.

Bound CHI3L1 was detected using a CHI3L1-specific primary antibody followed by an HRP-conjugated secondary antibody. Signal was developed using TMB substrate, stopped with sulfuric acid and measured at 450 nm. Values were normalized to the vehicle-treated condition for the corresponding CHI3L1 genotype. Concentration-response curves were fitted using four-parameter logistic regression. Three independent biological experiments were performed for each condition, with technical replicate ELISA measurements averaged within each experiment.

### ELISA-based signaling analysis

Cells were treated with vehicle, **CHI-DEL** or **CHI-DEL-Neg** for 6 h unless otherwise indicated. Cells were washed twice with ice-cold PBS and lysed in buffer supplemented with protease and phosphatase inhibitors. Lysates were clarified at 16,000 × *g* for 15 min at 4 °C, and total protein concentration was determined by BCA assay. Phospho-STAT3 (Tyr705) and total STAT3 were quantified using the STAT3 (pY705) + total STAT3 ELISA Kit (Abcam, ab126459). Phospho-NF-κB p65 (Ser536) and total NF-κB p65 were quantified using the RayBio Phospho-NF-κB p65 (Ser536) and Total NF-κB p65 ELISA Kit (RayBiotech, PEL-NFKBP65-S536). Assays were performed according to the manufacturers’ instructions using equivalent amounts of total protein. Phosphorylated protein signals were normalized to their corresponding total-protein signals and subsequently to vehicle within each genotype.

### Human iPSC-derived microglia

Human iPSCs were maintained on Matrigel-coated plates in mTeSR Plus medium and routinely confirmed to be free of mycoplasma contamination. Microglial differentiation was performed through a hematopoietic progenitor intermediate using a staged protocol adapted from established iPSC-microglia differentiation methods. Briefly, iPSCs were differentiated toward mesoderm and hematopoietic progenitor cells using BMP4, VEGF and SCF, followed by specification toward the myeloid lineage using M-CSF and IL-3 ^26,27^. Myeloid progenitors were matured for approximately 14 d in serum-free medium containing IL-34, M-CSF and TGF-β1. Final maturation was performed in the presence of IL-34, TGF-β1 and CX3CL1.

Independent differentiation batches were initiated separately and were treated as independent biological replicates. Before experimental use, microglial identity was verified by flow cytometry and immunophenotyping for CD11b, CD45, CX3CR1, P2RY12 and TMEM119. Cultures were required to contain at least 85% viable cells and at least 80% cells displaying the predefined microglial marker profile before inclusion in conditioned-medium experiments.

### Preparation of GBM-conditioned medium

Conditioned medium was generated independently from WT, CHI3L1-KO, KO+WT-rescue and KO+W99F-rescue GBM cells. Producer cells were seeded at matched density and allowed to equilibrate overnight. Cells were then washed twice with PBS and incubated in serum-reduced medium containing 1% fetal bovine serum for 24 h. Conditioned medium was collected and clarified by centrifugation at 1,000 × g for 10 min at 4 °C followed by centrifugation at 10,000 × g for 10 min to remove residual cellular material. CHI3L1 concentration in conditioned medium was quantified by ELISA to confirm loss of secretion in the knockout cells and comparable secretion in WT- and W99F-rescue cells.

Recipient iPSC-derived microglia were exposed to conditioned medium for 24 h. Where indicated, microglia were simultaneously treated with vehicle, 1 µM **CHI-DEL** or 1 µM **CHI-DEL-Neg**. The complete experiment comprised the nine treatment conditions shown in Figure 5.

### Microglial flow cytometry

Following conditioned-medium exposure, adherent microglia were gently detached using Accutase, washed in PBS containing 2% fetal bovine serum and incubated with Fc receptor blocking reagent for 10 min at 4 °C. Cells were stained with a fixable viability dye together with antibodies against CD45, CD11b, P2RY12 and CD206 for 30 min at 4 °C protected from light. Cells were washed twice and resuspended in flow cytometry buffer.

Samples were acquired on a BD LSRFortessa and analyzed using FlowJo v10. Sequential gating consisted of exclusion of debris by forward- and side-scatter, singlet selection using FSC-A versus FSC-H, exclusion of nonviable cells, identification of CD45⁺CD11b⁺P2RY12⁺ microglia, and quantification of CD206-positive cells within the resulting population. Single-color compensation controls were included in each experiment. Fluorescence-minus-one controls were used to define the CD206 positivity threshold. An identical gating template was applied to all samples from a given differentiation batch. Three independent iPSC-microglial differentiation batches were analyzed per experimental arm (n = 3); replicate wells within each differentiation batch were averaged before statistical analysis.

### Quantitative RT-PCR

Total RNA was isolated from treated iPSC-derived microglia using the RNeasy Mini Kit with on-column DNase digestion. RNA concentration and purity were determined spectrophotometrically, and equal amounts of RNA were reverse-transcribed using the High-Capacity cDNA Reverse Transcription Kit. Quantitative PCR was performed using PowerUp SYBR Green Master Mix on a QuantStudio real-time PCR instrument. Validated primer pairs were used for SPP1, ARG1 and GAPDH. Relative transcript abundance was calculated using the ΔCq method with GAPDH as the internal reference gene. Three independent iPSC-microglial differentiation batches were analyzed per experimental arm (n = 3). qPCR reactions were performed in technical triplicate, and technical wells were averaged to yield one value per differentiation batch before statistical analysis.

### IL-10 ELISA

Microglial culture supernatants were collected after 24 h of conditioned-medium exposure, clarified by centrifugation at 1,000 × *g* for 10 min at 4 °C, aliquoted, and stored at -80 °C until analysis. Human IL-10 concentrations were quantified using the Human IL-10 Quantikine ELISA Kit (R&D Systems/Bio-Techne, Cat. No. D1000B) according to the manufacturer’s instructions. Samples were thawed on ice and analyzed in duplicate together with recombinant human IL-10 standards. Absorbance was measured at 450 nm with wavelength correction at 540 nm. Background-corrected absorbance values were used to generate a standard curve by four-parameter logistic regression, and IL-10 concentrations were interpolated from the fitted curve. Four independent biological replicates were analyzed per experimental arm (n = 4), with duplicate ELISA measurements averaged to yield one value per biological replicate.

### Microglia-to-GBM feedback invasion assay

To determine whether CHI3L1-dependent microglial remodeling altered GBM invasive behavior, supernatants from treated iPSC-derived microglia were transferred to GBM cells in a secondary Matrigel invasion assay. GBM cells were serum-starved for 6 h and 5 × 10^4^ cells were seeded into the upper compartment of 8-µm Matrigel-coated Transwell inserts in serum-free medium. Microglial conditioned medium was added to the lower chamber as the sole conditioned stimulus. Cells were allowed to invade for 20 h. Noninvading cells were removed, and invaded cells were fixed in 4% paraformaldehyde and stained with crystal violet. Five fields per insert were quantified under identical imaging conditions. Technical values were averaged within each insert and replicate. Four independent biological experiments were performed per treatment arm.

### In vitro ADME and permeability profiling

The physicochemical and in vitro ADME properties of **CHI-DEL** were evaluated using standardized assays. LogD at pH 7.4 was determined by the shake-flask method using mutually saturated n-octanol and phosphate-buffered saline. Kinetic solubility was measured by diluting a 10 mM DMSO stock into phosphate-buffered saline to a final DMSO concentration of 1%, incubating the samples for 2 h at room temperature, and quantifying the soluble compound in the clarified supernatant by LC-MS/MS.

Passive permeability was measured using a parallel artificial membrane permeability assay (PAMPA). **CHI-DEL** was applied to the donor compartment at 10 µM and incubated for 4 h at room temperature. Compound concentrations in the donor and acceptor compartments were quantified by LC-MS/MS, and the apparent permeability coefficient (P_app_) was calculated from the rate of transfer across the artificial membrane. Bidirectional permeability was measured across confluent MDCK-MDR1 monolayers. **CHI-DEL** was applied at 5 µM to either the apical or basolateral compartment, and transport was measured over 120 min at 37 °C. Monolayer integrity was confirmed by transepithelial electrical resistance and transport of Lucifer yellow. The assay was performed in the absence or presence of 10 µM elacridar. Apparent permeability was calculated in the apical-to-basolateral and basolateral-to-apical directions, and the efflux ratio was calculated as P_app_,B→A/P_app_,A→B.

Microsomal stability was evaluated by incubating **CHI-DEL** at 1 µM with pooled mouse or human liver microsomes at a microsomal protein concentration of 0.5 mg ml^-1^ in the presence of 1 mM NADPH at 37 °C. Aliquots were collected at different time points and quenched with ice-cold acetonitrile containing an internal standard. Remaining parent compound was quantified by LC-MS/MS. The disappearance half-life was determined from the first-order depletion rate, and intrinsic clearance was calculated from the microsomal protein concentration and fitted elimination-rate constant. Plasma stability was evaluated by incubating **CHI-DEL** at 1 µM in pooled mouse or human plasma at 37°C. Samples were collected over 180 min, quenched with acetonitrile containing internal standard, and analyzed by LC-MS/MS. Human plasma protein binding was measured by equilibrium dialysis at a compound concentration of 2 µM after incubation for 4 h at 37 °C. hERG channel inhibition was evaluated by automated patch-clamp electrophysiology in hERG-expressing HEK293 cells across concentrations of 0.1-50 µM. Inhibition of CYP3A4, CYP2D6, CYP2C9, and CYP2C19 was measured at 10 µM using isoform-selective probe substrates in pooled human liver microsomes. All quantitative ADME measurements were performed in triplicate. The complete profile is provided in Table S2.

### Murine glioma cell lines

GL261 murine glioma cells were obtained from Creative Bioarray and maintained in DMEM supplemented with 10% fetal bovine serum, 1% GlutaMAX, and 1% penicillin-streptomycin. Cells were maintained at 37 °C in a humidified atmosphere containing 5% CO_2_. To generate GL261-CHI3L1 cells, parental GL261 cells were transduced with a lentiviral vector encoding human CHI3L1 under the control of the EF1α promoter. Stable cells were selected with 2 µg ml^-1^ puromycin, and a pooled population was used to minimize clone-specific effects.

### Orthotopic glioma model

All animal studies were performed in accordance with procedures approved by the Weill Cornell Medicine Institutional Animal Care and Use Committee (Protocol # 2024-0008).

Eight- to ten-week-old C57BL/6 mice were used for orthotopic GL261-CHI3L1 studies ^28^. Both sexes were included. Survival cohorts contained equal numbers of female and male mice, with five females and five males per treatment group. Mice were anesthetized with inhaled isoflurane and positioned in a stereotactic frame. A sterile guide screw was implanted over the predetermined intracranial injection site. After recovery from screw implantation, 1 × 10^5^ GL261-CHI3L1 cells suspended in 5 µl DPBS were injected intracranially through the guide screw at a controlled rate. Treatment began 5 d after tumor implantation. **CHI-DEL** or **CHI-DEL-Neg** was formulated in 5% DMSO, 10% Solutol HS15 and 85% sterile saline and administered intraperitoneally twice weekly for 3 weeks, corresponding to six total doses. For the GL261-CHI3L1 survival study, mice were assigned to vehicle, **CHI-DEL** (10 mg kg^-1^), **CHI-DEL** (20 mg kg^-1^), **CHI-DEL** (40 mg kg^- 1^) or **CHI-DEL-Neg** (40 mg kg^-1^) groups.

Animals were followed for up to 60 d after implantation or until a predefined humane endpoint was reached. Humane endpoints included greater than 20% loss of baseline body weight, body-condition score of 2/5 or lower, severe neurological impairment, inability to ambulate or access food or water, persistent recumbency, or other evidence of substantial distress requiring euthanasia. Animals were randomized to treatment groups before the first dose using a computer-generated sequence while balancing sex across groups. Drug formulations were coded so that personnel performing routine clinical scoring and terminal sample analysis were blinded to treatment identity. No animal was excluded from analysis after treatment allocation unless a prespecified technical exclusion unrelated to therapeutic response occurred.

### Pharmacokinetic analysis

Pharmacokinetics of **CHI-DEL** were evaluated in tumor-bearing C57BL/6 mice following a single intraperitoneal dose of 20 mg kg^-1^. Plasma, brain and intracranial tumor samples were collected at 0.25, 0.5, 1, 2, 4, 8 and 24 h after dosing. Three mice were sampled at each time point. Blood was collected into K₂EDTA tubes and centrifuged at 2,000 × g for 10 min at 4 °C to obtain plasma.

Brain and tumor tissues were weighed and homogenized at a 1:4 tissue-to-buffer ratio in ice-cold 50% methanol/water using a bead-based tissue homogenizer. **CHI-DEL** concentrations were quantified by LC-MS/MS using a triple-quadrupole mass spectrometer coupled to UHPLC. Chromatographic separation was performed on a C18 reversed-phase column using a water/acetonitrile gradient containing 0.1% formic acid. Analyte detection was performed in positive electrospray-ionization mode using multiple-reaction monitoring. A structurally related internal standard was added to plasma, brain and tumor samples before protein precipitation. Plasma samples were precipitated with three volumes of acetonitrile containing internal standard. Homogenized brain and tumor samples were extracted using the same solvent ratio. Samples were vortexed, centrifuged at 16,000 × g for 10 min, and the resulting supernatant was subjected to LC-MS/MS.

Matrix-matched calibration curves were prepared separately in plasma, brain homogenate and tumor homogenate. Calibration curves covered 1-5,000 ng ml^-1^ or the corresponding tissue concentration range and were fitted using weighted linear regression. The lower limit of quantification was 1 ng ml⁻¹. Quality-control samples at low, medium and high concentrations were required to fall within ±15% of nominal concentration, or ± 20% at the LLOQ. Noncompartmental pharmacokinetic analysis was performed using Phoenix WinNonlin. C_max_ and T_max_ were taken directly from the observed concentration-time profiles. AUC was calculated using the linear-up/log-down trapezoidal method, and terminal half-life was estimated from the terminal log-linear phase where at least three quantifiable points were available. Brain-to-plasma and tumor-to-plasma concentration ratios were calculated at matched time points.

### Tumor target engagement and pharmacodynamic analysis

Separate cohorts of mice bearing orthotopic GL261-CHI3L1 tumors received vehicle or **CHI-DEL** at 10, 20 or 40 mg kg⁻¹ using the twice-weekly treatment schedule (n = 8 mice per dose). Tumors were collected 4 h after the final scheduled dose to align terminal pharmacodynamic measurements with a period of measurable tumor exposure. Mice were euthanized, tumors were rapidly dissected on ice and divided into portions for CETSA, CHI3L1-CD44 complex analysis and signaling measurements.

For tumor CETSA, approximately 30 mg tumor tissue was homogenized in ice-cold PBS containing protease inhibitors. Homogenates were divided into equal aliquots and heated at 55 °C for 3 min, followed by 3 min at room temperature. Samples were lysed by three freeze-thaw cycles and centrifuged at 20,000 × g for 20 min at 4 °C. Soluble CHI3L1 in the supernatant was quantified by ELISA and normalized to total protein in the corresponding homogenate. Values were subsequently expressed relative to the vehicle group. CHI3L1-CD44 complex abundance was measured using the plate-based complex assay described above and normalized to vehicle-treated tumors. Phospho-STAT3 signal was normalized to total STAT3 and subsequently to the vehicle group.

### Tumor immune profiling

Tumors from mice treated with vehicle, **CHI-DEL** at 20 mg kg^-1^ or **CHI-DEL-Neg** at 20 mg kg^-1^ (n = 8 mice per group) were collected 4 h after the final scheduled dose. Fresh tumor tissue was minced mechanically and enzymatically dissociated for 30 min at 37 °C in RPMI containing collagenase D and DNase I with gentle agitation. Digests were passed through 70-µm cell strainers, washed and subjected to red blood cell lysis when required. Single-cell suspensions were stained with fixable viability dye followed by antibodies against CD45, CD11b, F4/80, CD3, CD8, CD206 and SPP1. For intracellular SPP1 analysis, cells were fixed and permeabilized after surface staining. Samples were acquired on a BD LSRFortessa and analyzed using FlowJo v10.

Sequential gating consisted of forward/side-scatter selection, singlet discrimination and exclusion of nonviable cells. Tumor-associated macrophages were defined as viable CD45⁺CD11b⁺F4/80⁺ cells. CD206-positive and SPP1-positive fractions were quantified within the TAM population. CD8⁺ T cells were defined as viable CD45⁺CD3⁺CD8⁺ cells and were reported as a fraction of total CD45⁺ immune cells. Single-color compensation controls and fluorescence-minus-one controls were used for compensation and gate placement. The same gating hierarchy and positivity thresholds were applied across all treatment conditions within an experiment. Each plotted point represents one animal.

### Animal tolerability

Body weight and general clinical condition were assessed at least three times weekly during treatment and daily when animals approached predefined humane endpoints. Body-weight change was calculated relative to the pretreatment value obtained immediately before the first compound dose. Clinical observations included posture, grooming, mobility, neurological behavior, hydration, food intake and body-condition score. Injection-site reactions and other overt signs of compound-related toxicity were recorded throughout the study.

Repeat-dose tolerability was evaluated in non-tumor-bearing C57BL/6 mice assigned to vehicle or **CHI-DEL** at 10, 20, or 40 mg kg^-1^ (n = 3 mice per group). Treatments were administered intraperitoneally twice weekly for 3 weeks, corresponding to six total doses, using the same formulation as in the efficacy studies. Body weight and general clinical condition were recorded before treatment and at least three times weekly throughout the 21-day observation period.

### Statistical analysis

Statistical analyses were performed using GraphPad Prism 10 and R 4.5.0. Sample sizes were selected on the basis of prior experience with the corresponding cellular and orthotopic glioma models and were not adjusted on the basis of interim treatment effects. Biological and technical replication were distinguished throughout the study. Technical replicate wells or measurements were averaged before inferential testing. Statistical n denotes independent biological experiments, independent iPSC differentiation batches, independent protein preparations or individual animals, as appropriate. Data are presented as mean ± SEM unless otherwise stated. Individual biological observations are shown wherever practical. Two-group comparisons were performed using two-sided Welch’s t-tests unless another analysis is specified. For mutant-qualification experiments comparing multiple variants with WT, P values were adjusted using the Benjamini– Hochberg false-discovery-rate procedure. Concentration-response data were fitted by nonlinear regression using a four-parameter logistic model with variable slope.

Survival distributions were estimated by the Kaplan-Meier method and compared using two-sided log-rank tests. For the GL261 dose-ranging study, pairwise comparisons of active treatment groups against vehicle were adjusted using the Holm method. Comparisons between dose-matched **CHI-DEL** and **CHI-DEL-Neg** were treated as predefined pharmacological-specificity comparisons. Hazard ratios and 95% confidence intervals were calculated using Cox proportional-hazards regression as secondary measures of treatment effect. Animals alive at the predetermined end of follow-up were right-censored at that time. No responder/nonresponder classification was used for the primary survival analysis. P < 0.05 was considered statistically significant.

## Data availability

The atomic coordinates and structure factors for the **CHI-DEL** complex have been deposited in the Protein Data Bank under accession code 10JV. All other data supporting the findings of this study are available within the Article and its Supplementary Information. The human CHI3L1 amino acid sequence is available in UniProt under accession P36222. The additional PDB structures used for structural comparison are 1HJX, 1HJW, and 8R41.

## Author Contribution declaration

S.U. contributed to conceptualization, methodology, investigation, data analysis, structural and crystallographic analysis, visualization and preparation of the original manuscript draft. L.Z. contributed to methodology, investigation and data analysis. M.T. contributed to structural studies, crystallographic analysis and interpretation. S.K. contributed to investigation and data analysis. M.T.G. conceived and supervised the study, administered the project and reviewed and edited the manuscript. All authors reviewed and approved the final manuscript.

## Supporting Information

The Supporting Information is available free of charge on the ACS Publications website.

SPR sensorgrams, Chemical structure of **CHI-DEL-Neg**, Electrostatic surface representation of the CHI3L1-**CHI-DEL** complex, Full melt curves and intracellular CHI3L1 levels for the mutant qualification panel, validation of the isogenic CHI3L1 knockout/rescue panel, Brain and tumor tissue penetration, In vitro PK/ADME profile for **CHI-DEL**, Crystallographic data collection, refinement and model validation, NMR spectra, HPLC traces, and LC-MS data (PDF).

