## Supporting Information for "Resistance-Mutant Rescue Establishes On-Target CHI3L1 Inhibition by CHI-DEL, a Brain-Penetrant Small Molecule with In Vivo Efficacy in Glioblastoma"

**Structure-Guided Chemical Genetics Establishes On-Target Pharmacology for CHI3L1**

Moustafa Gabr^1,2*^

^1^Department of Radiology, Columbia University Irving Medical Center, New York, NY, 10032, USA

^2^Department of Radiology, Weill Cornell Medicine, New York, NY 10065, USA

^3^Structural Biology Program, Memorial Sloan Kettering Cancer Center, New York, NY 10065, USA

^4^International Health Division, Indian Council of Medical Research, Ansari Nagar, New Delhi 110029, India

* To whom correspondence should be addressed:

Moustafa T. Gabr.

**DNA-encoded library screening and candidate prioritization**

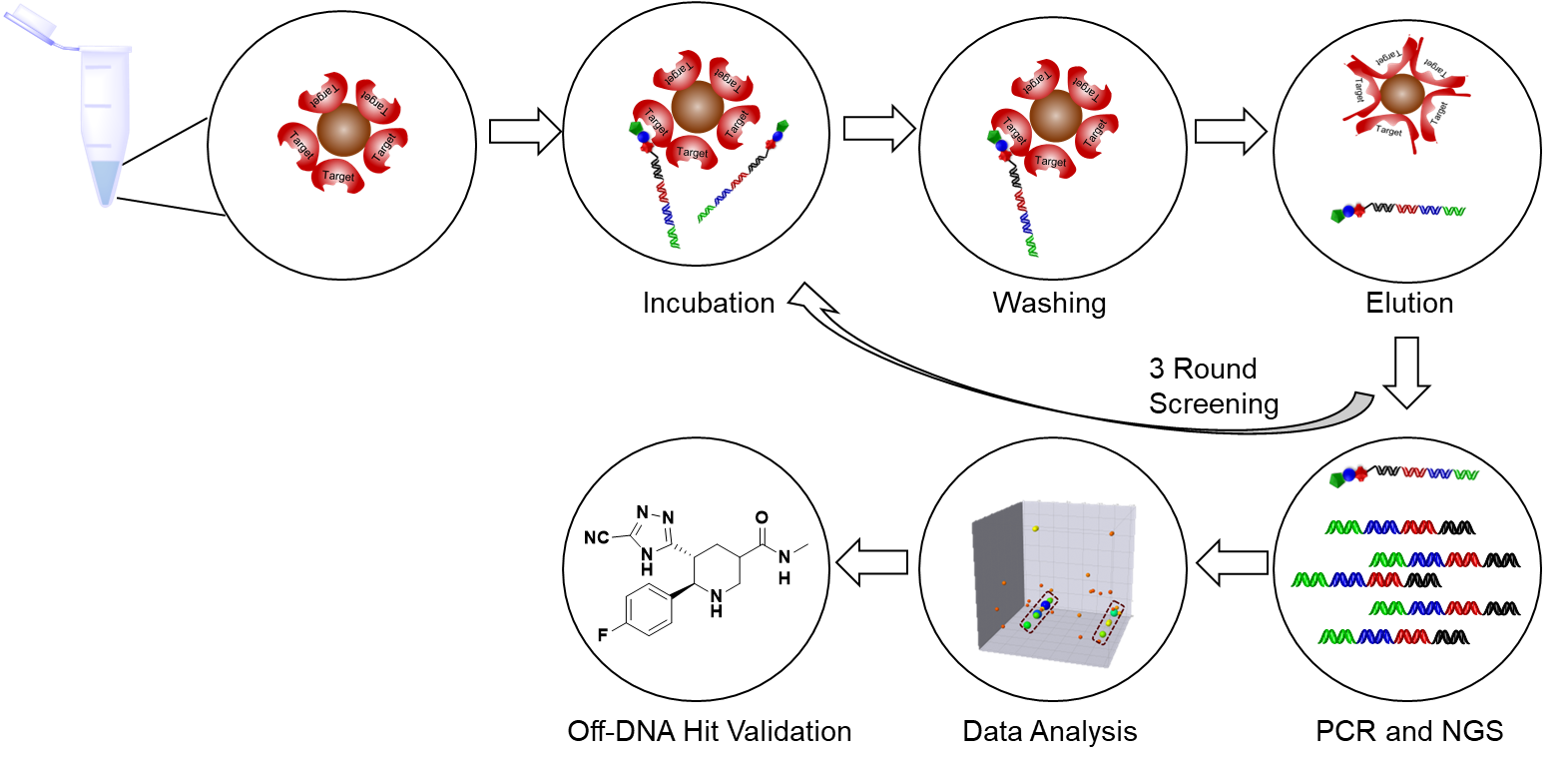

**Fig. S1. DNA-encoded library screening workflow for CHI3L1.** A 1.4-billion-member DNA-encoded library was incubated with recombinant human CHI3L1 immobilized on affinity beads, with a no-target bead control processed in parallel. Following washing to remove unbound library members, retained compounds were recovered and their associated DNA barcodes amplified for the subsequent selection round. After three consecutive rounds of affinity selection, recovered barcodes were analyzed by next-generation sequencing and decoded to identify enriched building-block combinations for off-DNA synthesis and biophysical validation.

**SPR evaluation of additional DEL-derived candidates**

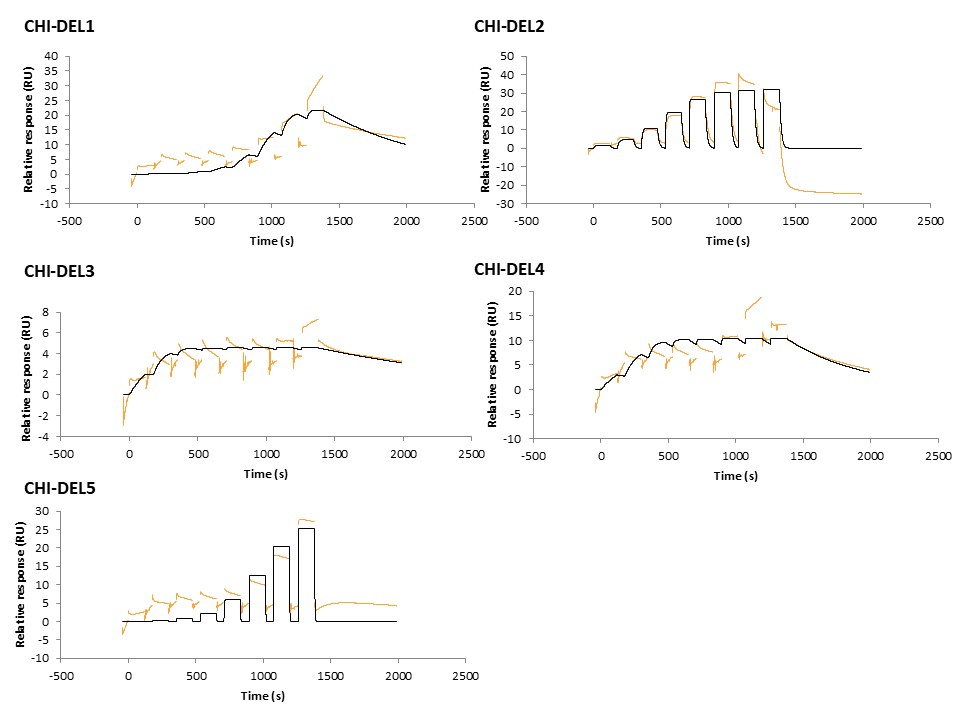

**Fig. S2. Sensorgram obtained during the initial SPR evaluation of Cpd A against immobilized recombinant human CHI3L1.** Experimental response is shown in orange, and the corresponding fit to a 1:1 binding model is shown in black. Despite showing the highest DNA-encoded library enrichment of all candidates prioritized against CHI3L1 (302.4-fold over the mean), the response amplitude was low (~4.5 RU maximum) across the tested concentration range, precluding reliable determination of affinity and kinetic parameters. **Cpd A** was not advanced for further characterization.

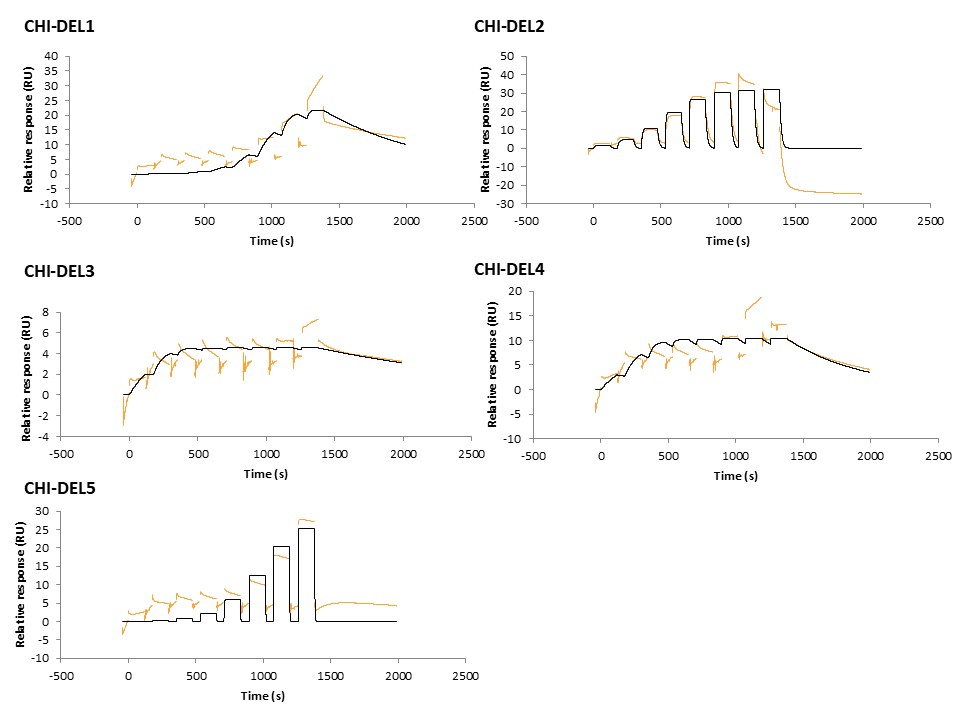

**Fig. S3. Sensorgram obtained during the initial SPR evaluation of Cpd B against immobilized recombinant human CHI3L1.** Experimental response is shown in orange, and the corresponding fit to a 1:1 binding model is shown in black. Although the response reached a higher maximum amplitude (~10 RU) than **Cpd A**, the experimental trace showed large, irregular excursions not captured by the fitted model, particularly during later injection cycles, precluding reliable determination of affinity and kinetic parameters. **Cpd B** was not advanced for further characterization.

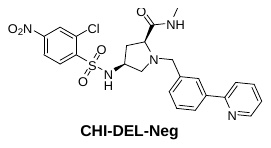

**Fig. S4. Chemical structure of CHI-DEL-Neg.**

**Electrostatic surface representation of the CHI3L1-CHI-DEL binding cleft.**

Electrostatic surface analysis further illustrates the position of **CHI-DEL** within the central CHI3L1 cleft (Supplementary Fig. S4). The ligand occupies a recessed region surrounded by negatively charged, positively charged and comparatively neutral surface patches. This chemically heterogeneous environment is consistent with the crystallographically observed binding mode, which includes nonpolar contacts together with geometrically plausible polar interactions involving Tyr206, Arg263 and Glu290. The surface representation further illustrates the accommodation of **CHI-DEL** between opposing sides of the cleft. Considered together with the residue-level structural analysis, these observations indicate that **CHI-DEL** recognition involves contacts distributed throughout the binding cavity rather than being dominated by a single localized electrostatic interaction.

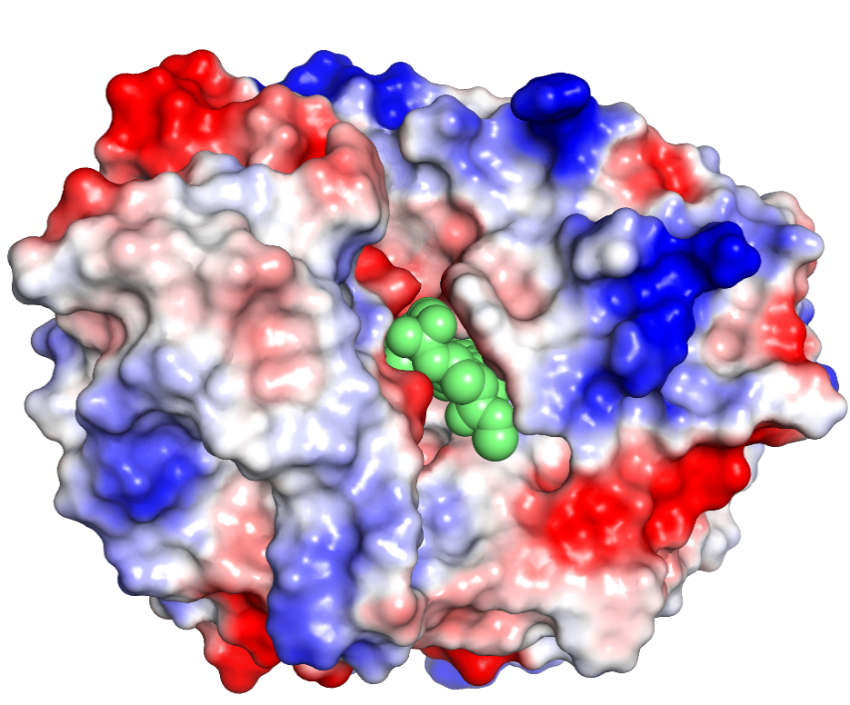

**Fig. S5. Electrostatic surface representation of the CHI3L1–CHI-DEL complex.** The solvent-accessible molecular surface of CHI3L1 is colored according to electrostatic potential, with negatively charged, positively charged and approximately neutral regions shown in red, blue and white, respectively. **CHI-DEL** is shown as green spheres within the central carbohydrate-binding cleft. The surface representation illustrates the recessed position of CHI-DEL and its accommodation within a chemically heterogeneous binding environment.

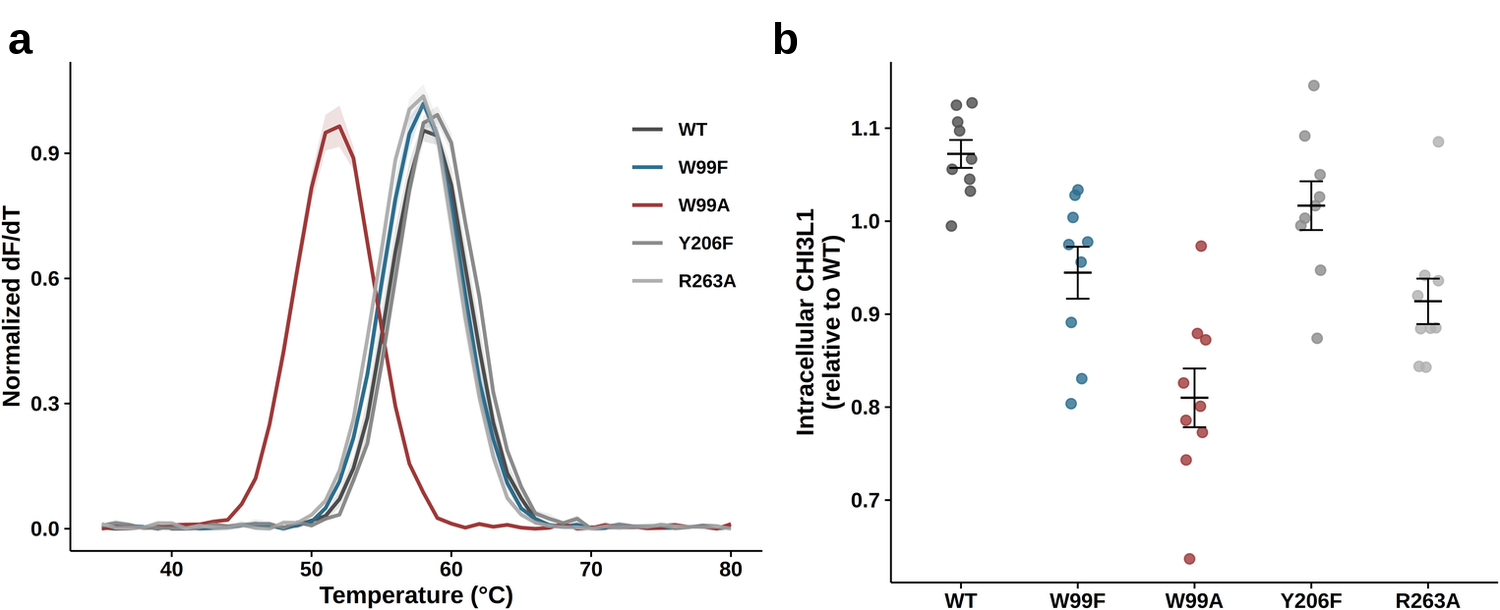

**Fig. S6. Full melt curves and intracellular CHI3L1 levels for the mutant qualification panel. (a)** Raw differential scanning fluorimetry (DSF) melt curves for all five variants (WT, W99F, W99A, Y206F, R263A). W99A shows a clear leftward shift (destabilization) consistent with the ~7°C Tm reduction reported in Figure 2; W99F, Y206F, and R263A overlap closely with WT. **(b)** Intracellular CHI3L1 level (relative to WT) for the same five variants. W99F shows a modest reduction relative to WT (mean ≈ 0.95 vs. 1.07), smaller than W99A's reduction (≈ 0.81) but non-negligible; Y206F and R263A are closer to WT. Data are mean ± SEM; each point is one biological replicate (n = 9/variant).

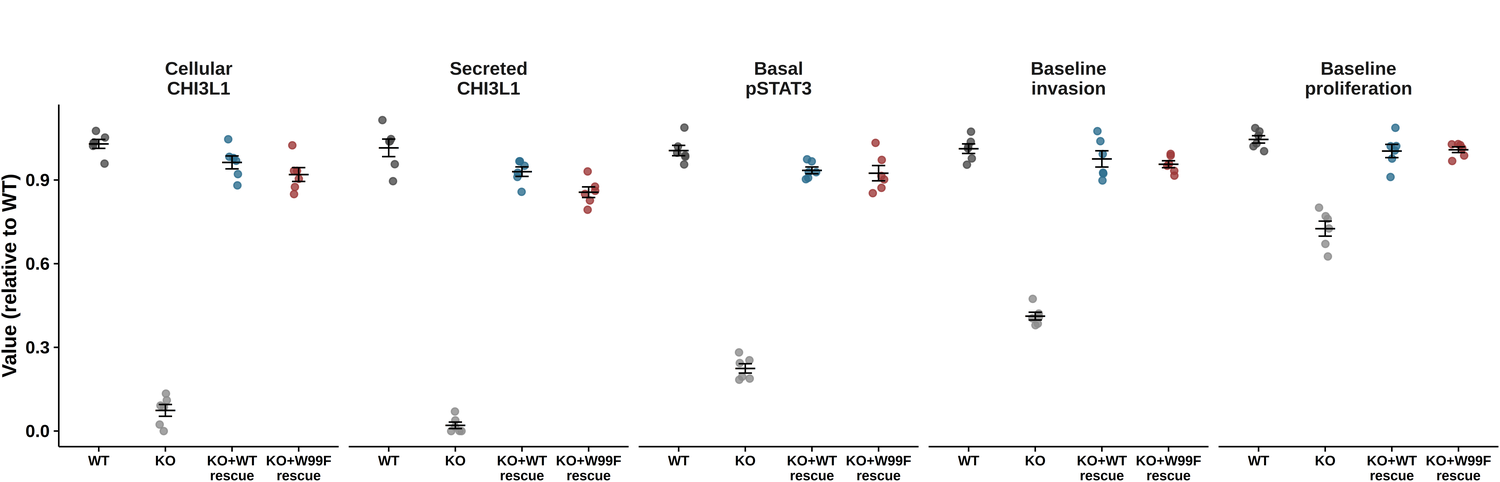

**Fig. S7. Baseline validation of the isogenic CHI3L1 knockout/rescue panel.** Cellular and secreted CHI3L1 level, basal pSTAT3 signaling, and baseline invasion and proliferation (all relative to WT, prior to any compound treatment) across WT, CHI3L1-KO, KO+WT-rescue, and KO+W99F-rescue cells. CHI3L1-KO reduces all five readouts relative to WT; both WT-rescue and W99F-rescue restore each readout to a comparable degree, confirming the two rescue lines are baseline-matched before any **CHI-DEL** treatment is introduced in Figures 3-5. Data are mean ± SEM; each point is one biological replicate (n = 6/genotype).

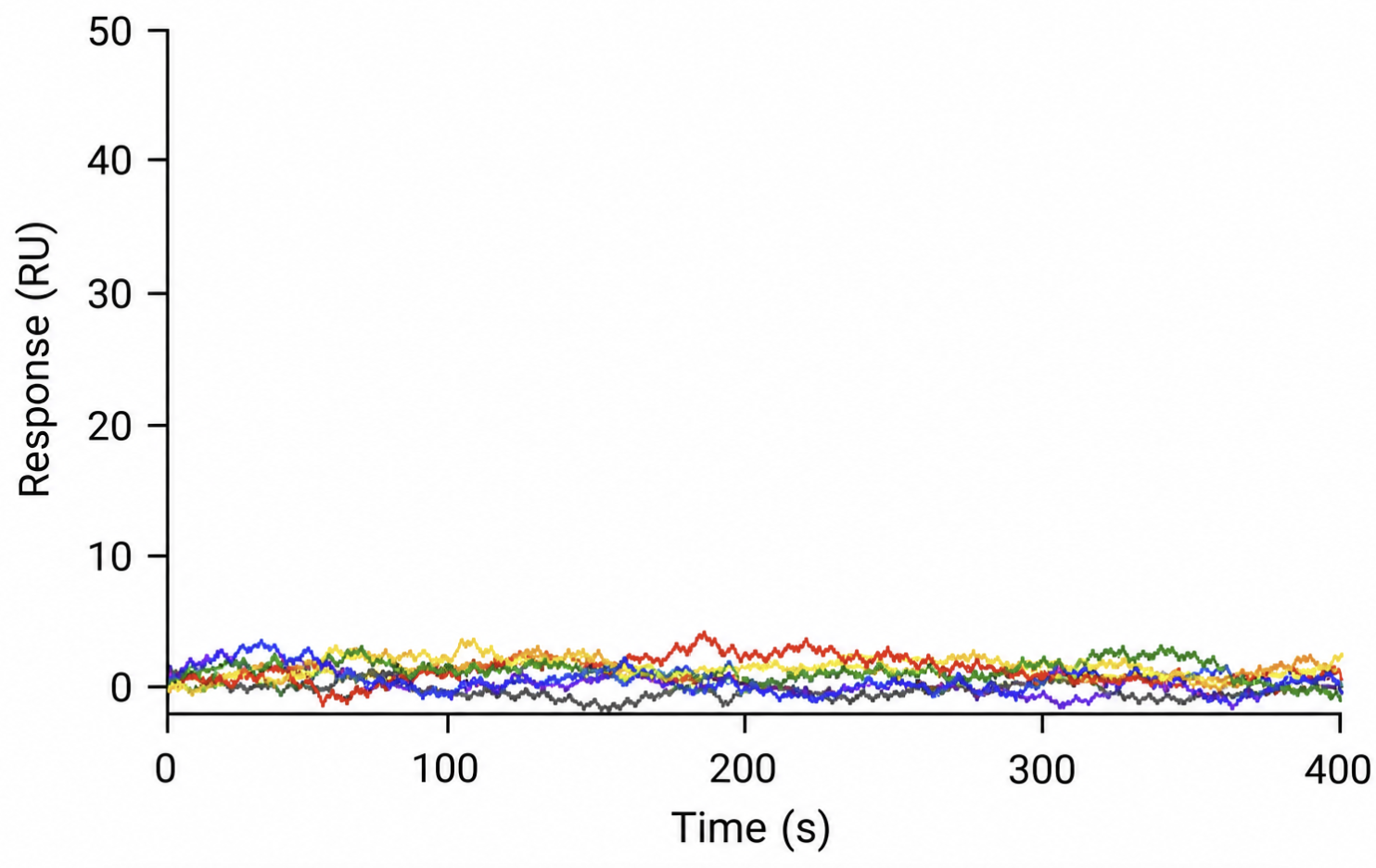

**Fig. S8.** Representative SPR sensorgram for **CHI-DEL** against purified W99F CHI3L1 across a concentration series up to 5 µM, showing no concentration-ordered, saturable response.

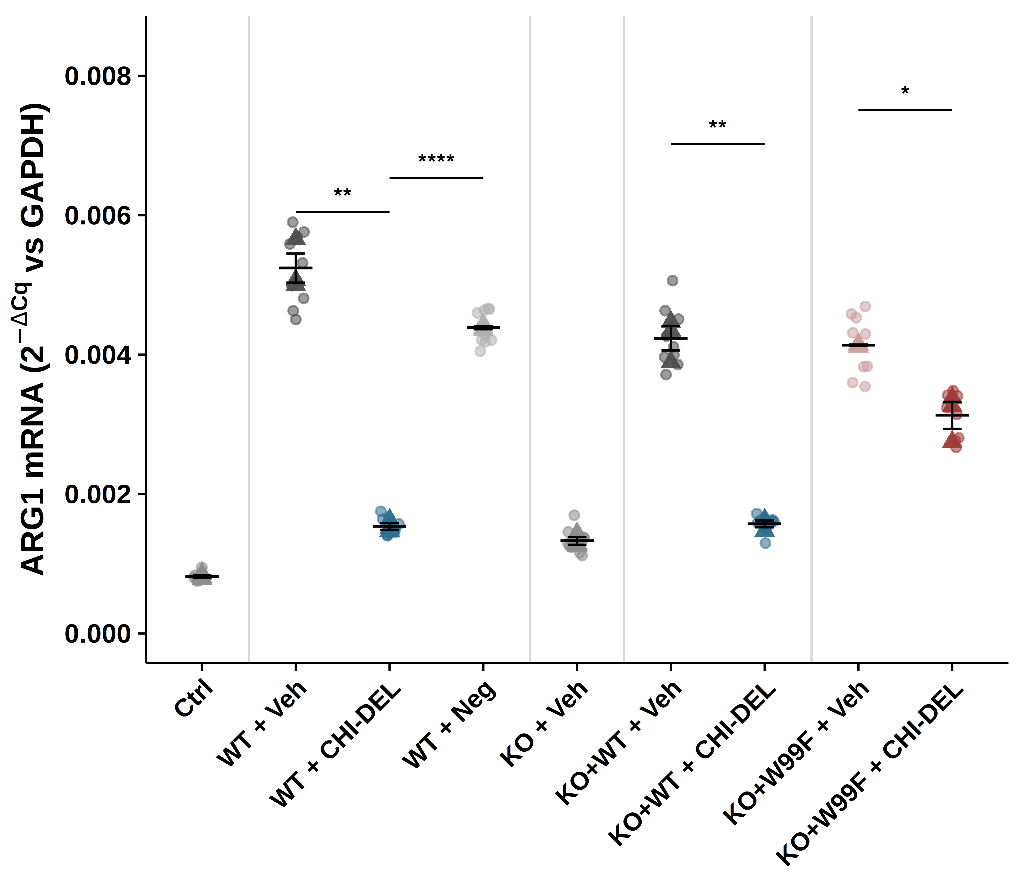

**Fig. S9. ARG1 mRNA expression in iPSC-derived microglia exposed to CHI3L1-genotyped GBM conditioned medium.** ARG1 mRNA expression (2^−ΔCq^ relative to GAPDH) by qPCR in the same recipient microglia and qPCR runs described in Figure 5c. Small points show individual replicate wells; triangles show the mean of each independent qPCR run (n = 3 runs/arm), the unit used for statistical testing. Thin vertical lines demarcate genotype blocks. Data are mean ± SEM. Brackets show two-sided Welch's *t*-tests for the same four mechanism-defining comparisons as Figure 5 (WT+Vehicle vs. WT+**CHI-DEL**, *t* = 17.3, *p* = 0.0021; WT+**CHI-DEL** vs. WT+**CHI-DEL-Neg**, *t* = 53.8, *p* < 0.0001; KO+WT-rescue Vehicle vs. **CHI-DEL**, *t* = 14.6, *p* = 0.0025; KO+W99F-rescue Vehicle vs. CHI-DEL, *t* = 5.17, *p* = 0.035). All comparisons significant; the KO+W99F-rescue effect is attenuated relative to KO+WT-rescue but not abolished, consistent with the pattern observed for CD206 and SPP1 in Figure 5.

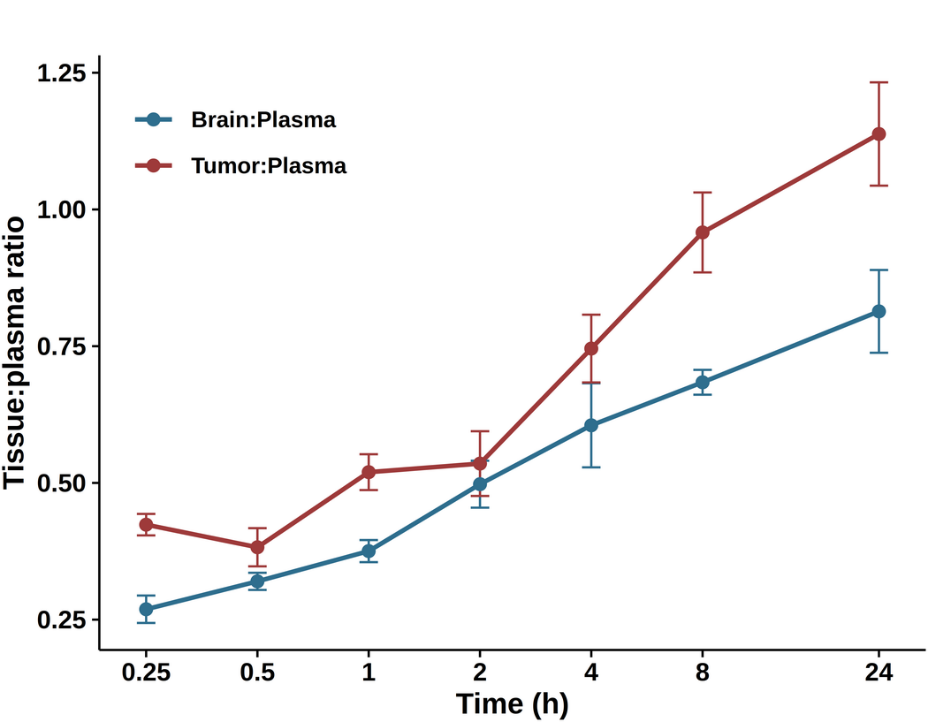

**Fig. S10. Brain and tumor tissue penetration ratios over time.** Brain:plasma and tumor:plasma concentration ratios following a single 20 mg/kg dose of **CHI-DEL**, calculated from the same animals shown in Figure 6a. Both ratios increase progressively over the 24 h profile, with tumor:plasma exceeding 1.0 by 24 h, consistent with tissue retention as plasma concentration declines faster than tissue concentration. Data are mean ± SEM (n = 3 mice per time point).

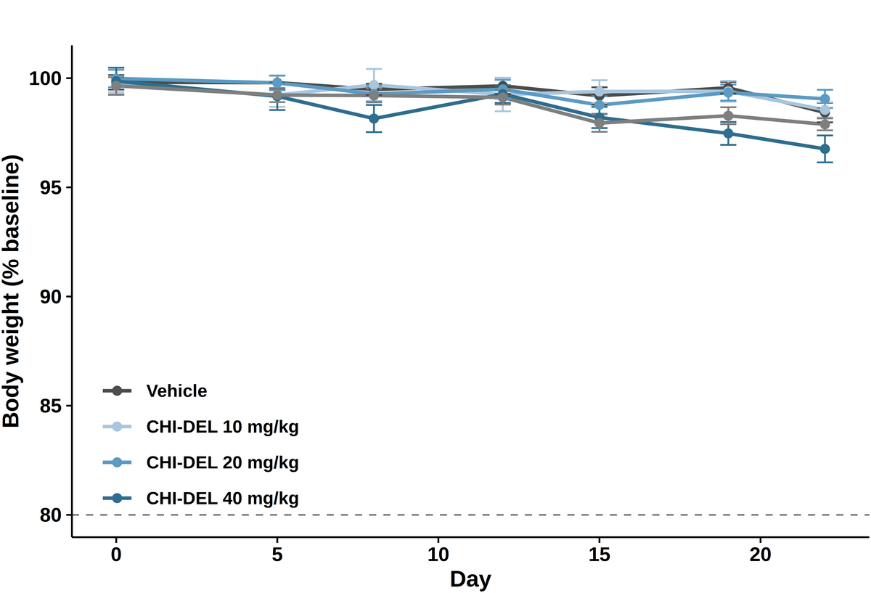

**Fig. S11. Body weight during repeat dosing.** Body weight (% of baseline) over 21 days of twice-weekly dosing with vehicle or CHI-DEL (10, 20, or 40 mg/kg) in non-tumor-bearing mice. No group showed weight loss approaching the 80% humane-endpoint threshold (dashed line) at any timepoint, including the highest dose tested. Data are mean ± SEM (n = 3 mice per group).

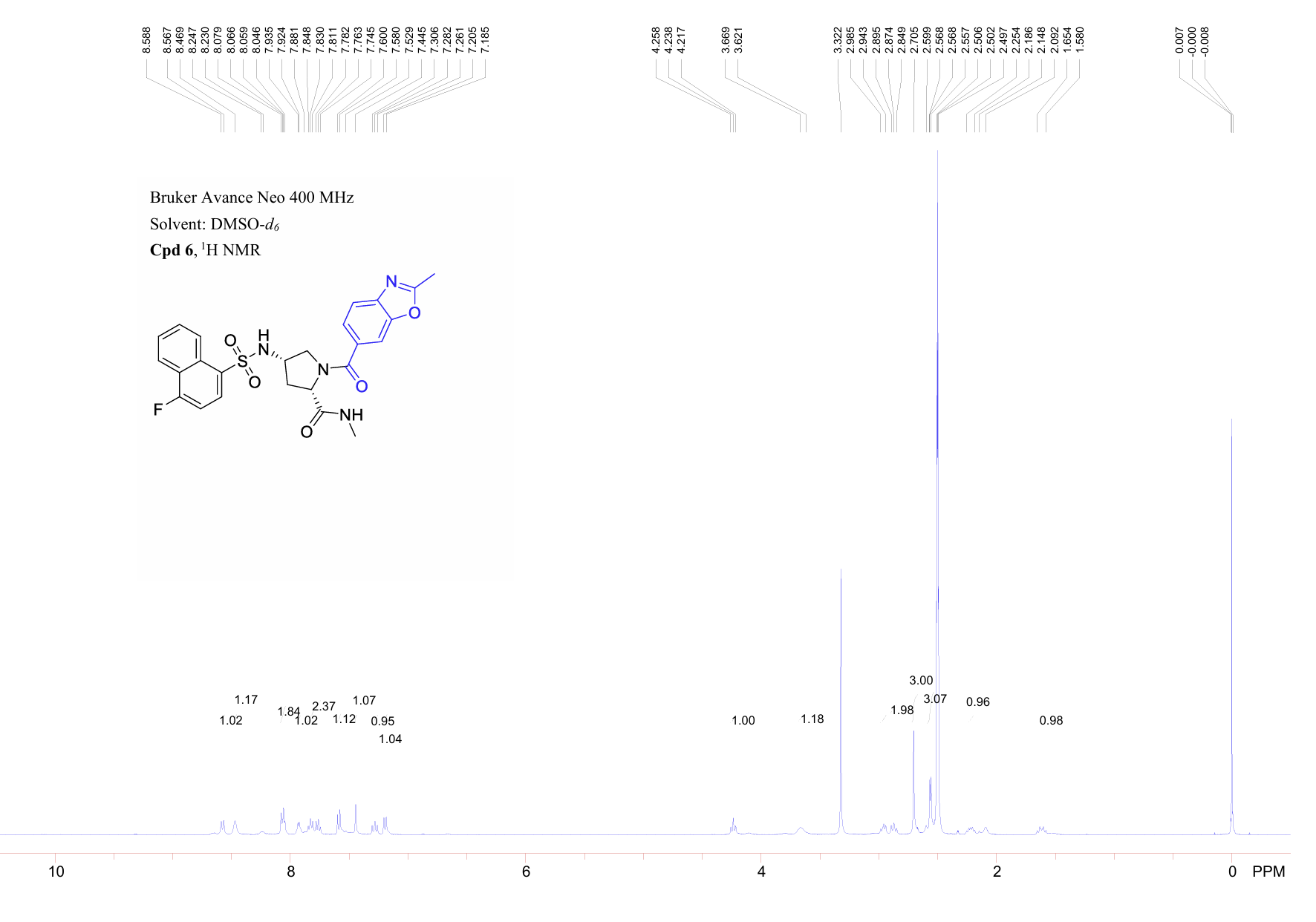

**Fig. S12.** ^1^H NMR spectrum for **CHI-DEL**.

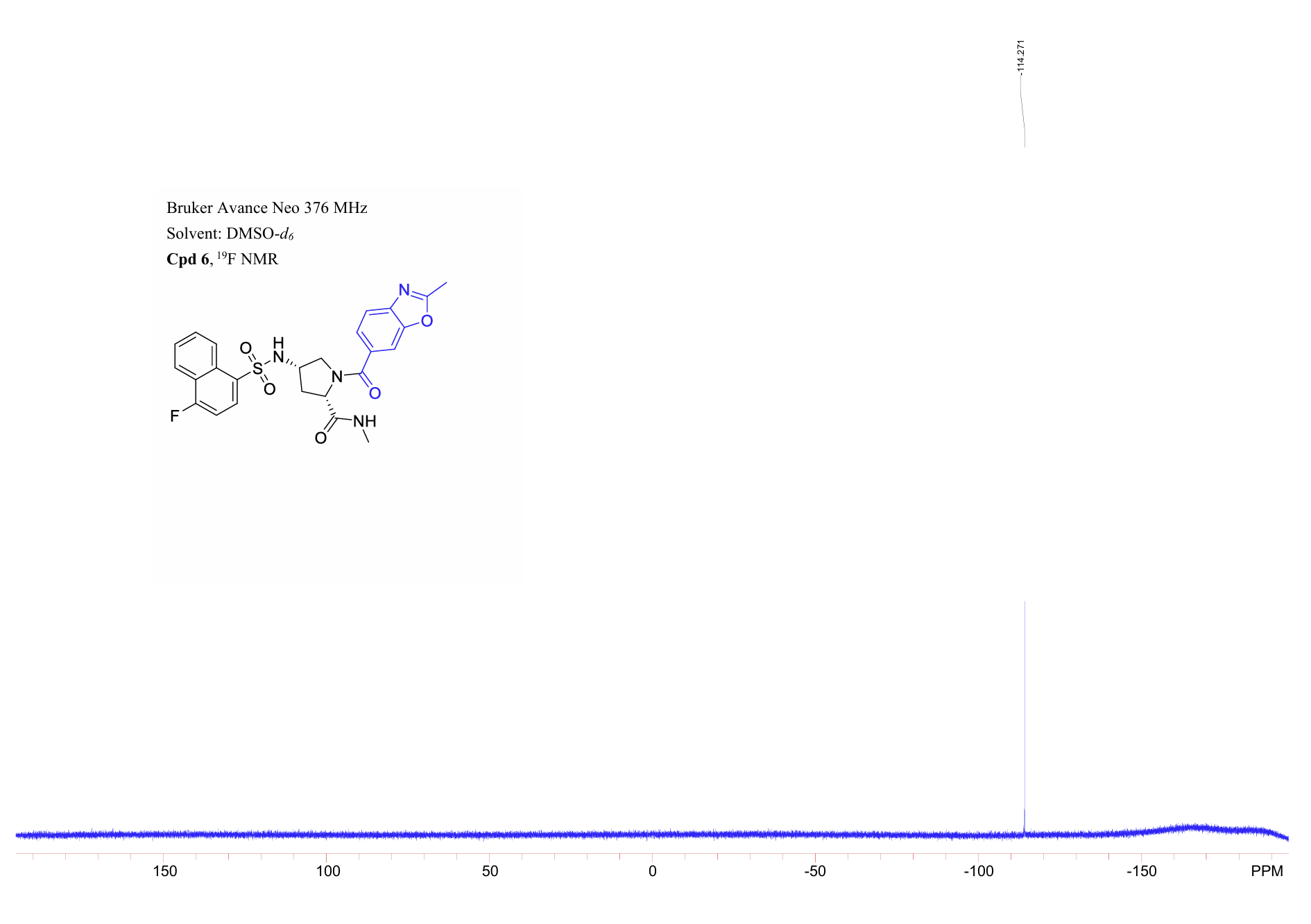

**Fig. S13.** ^19^F NMR spectrum for **CHI-DEL**.

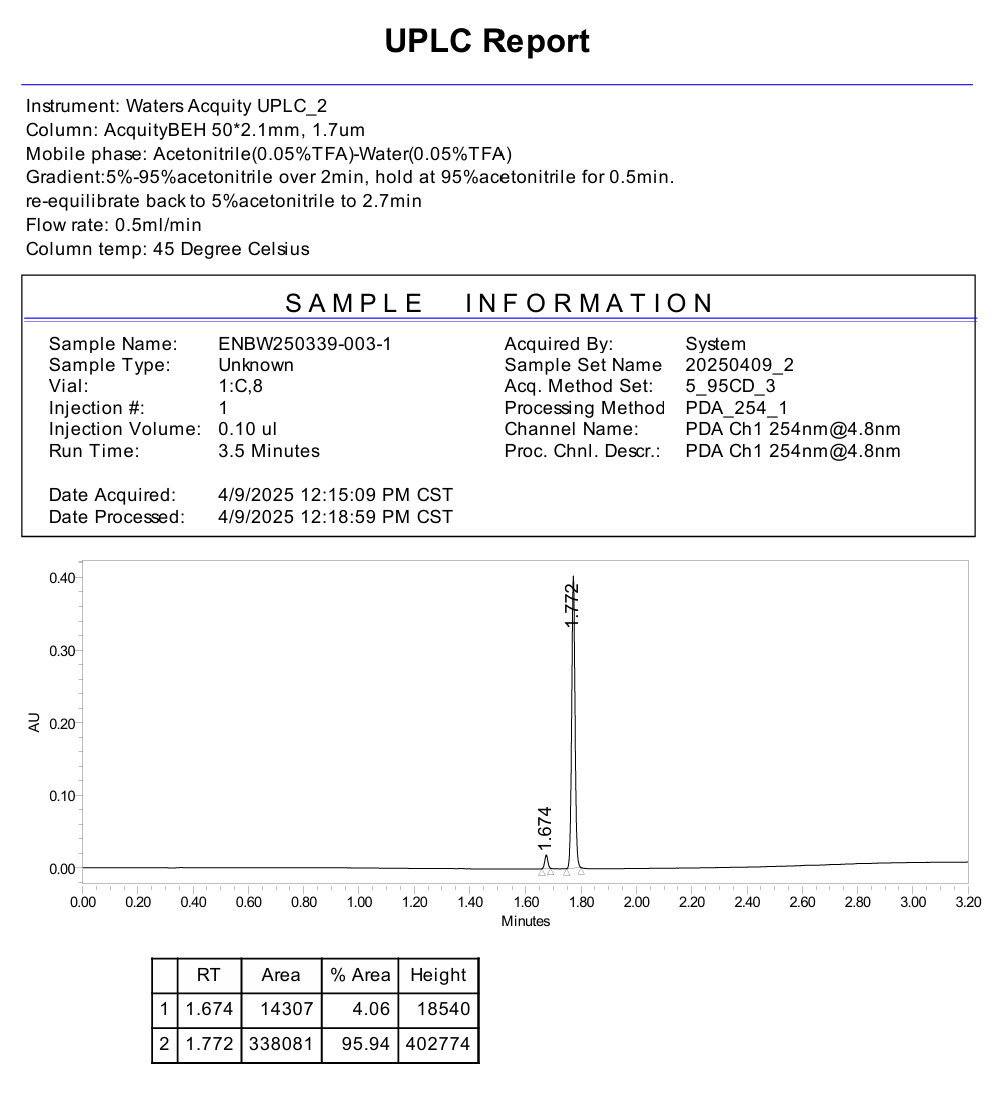

**Fig. S14.** UPLC purity report for **CHI-DEL**.

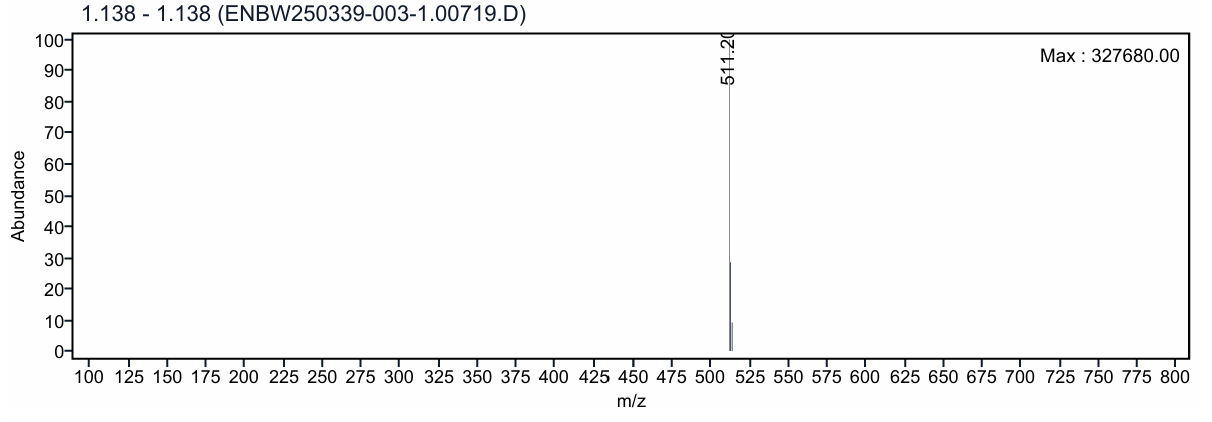

**Fig. S15.**  LCMS for **CHI-DEL**.

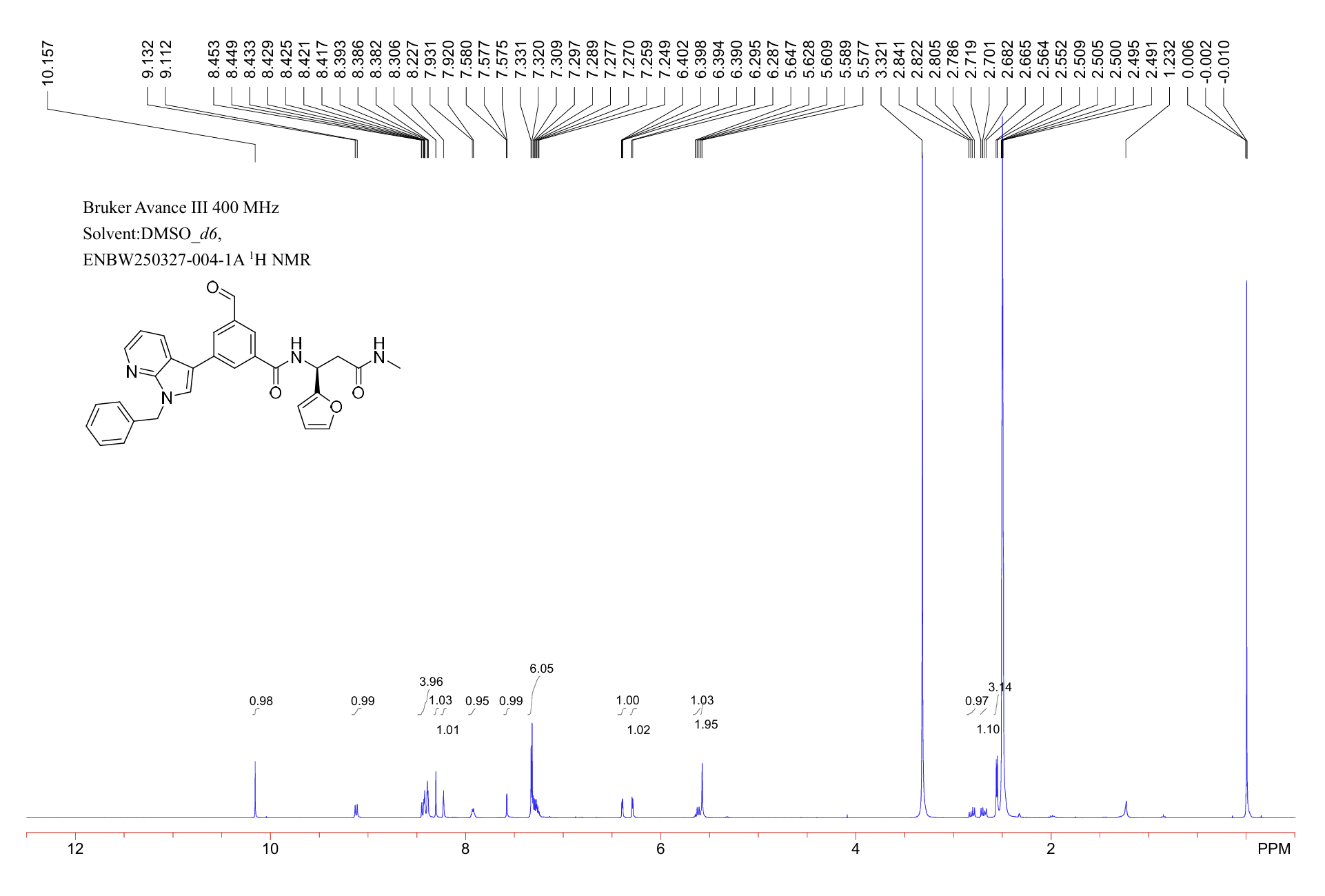

**Fig. S16.** ^1^H NMR spectrum for **Cpd A**.

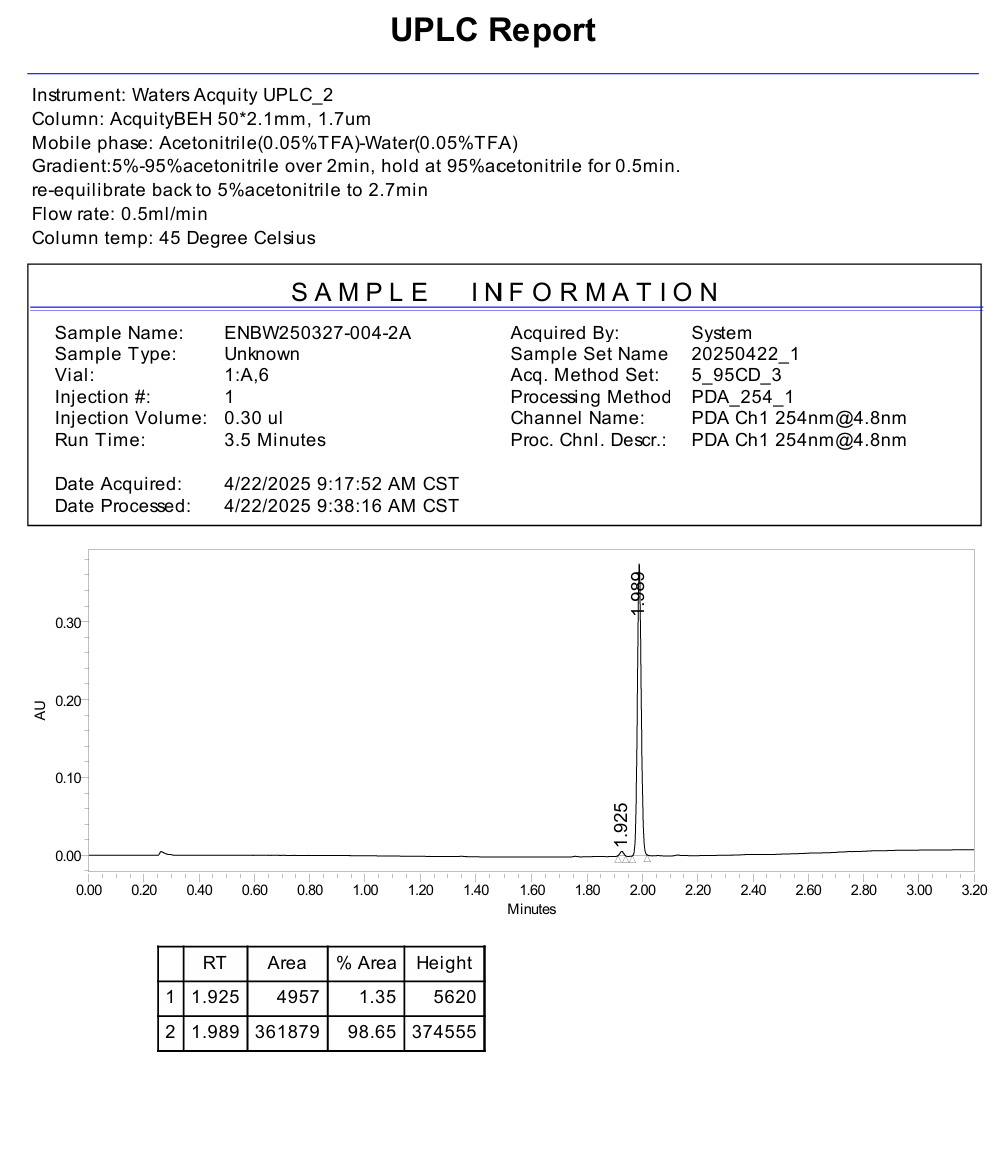

**Fig. S17.** UPLC purity report for **Cpd A**.

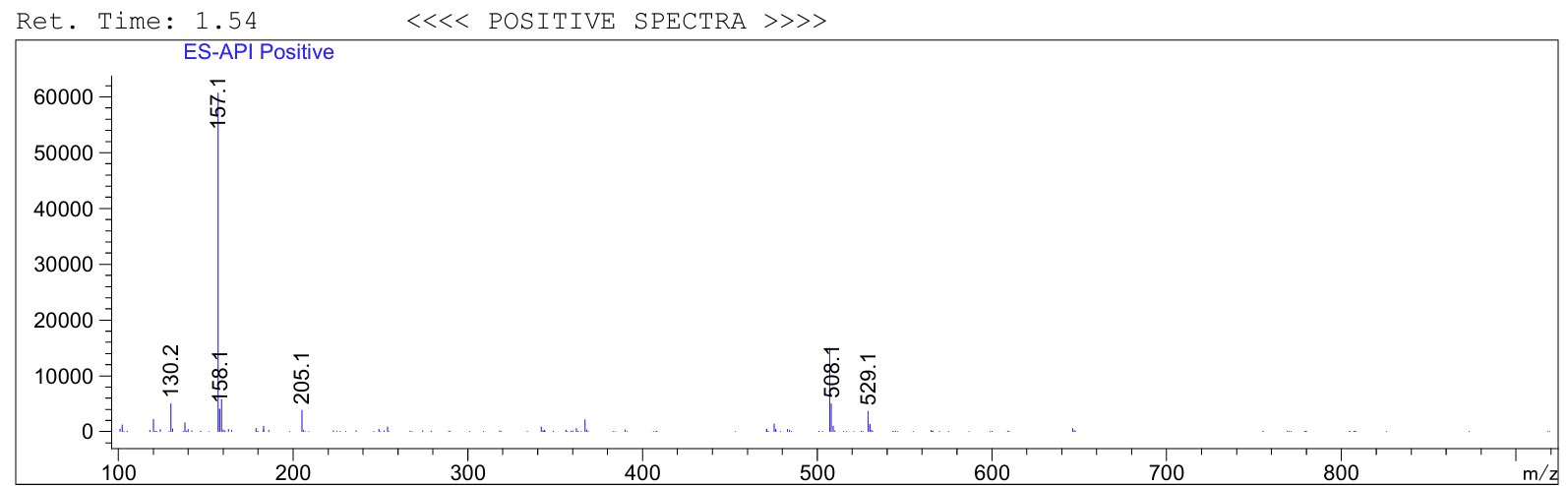

**Fig. S18.**  LCMS for **Cpd A**.

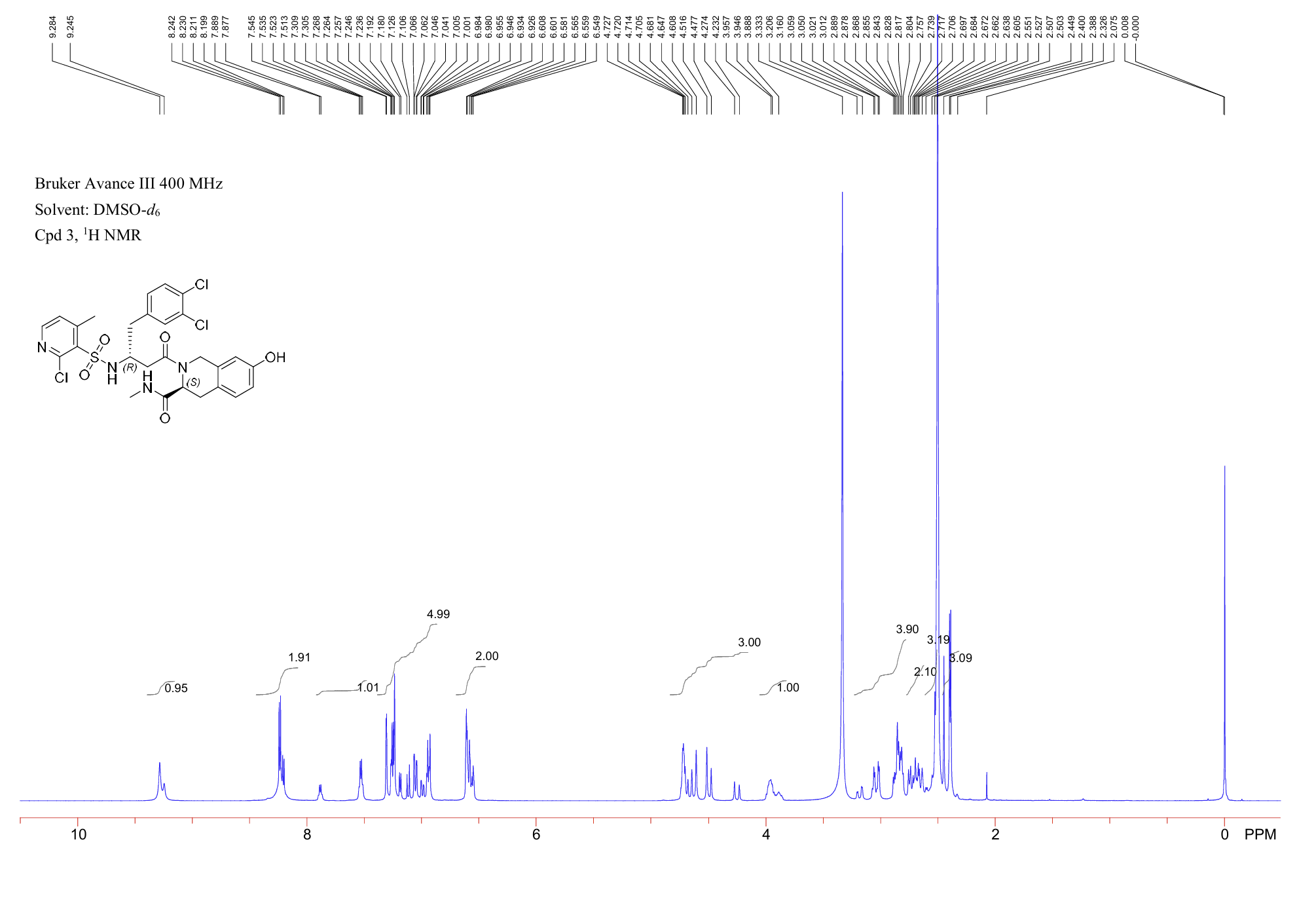

**Fig. S19.** ^1^H NMR spectrum for **Cpd B**.

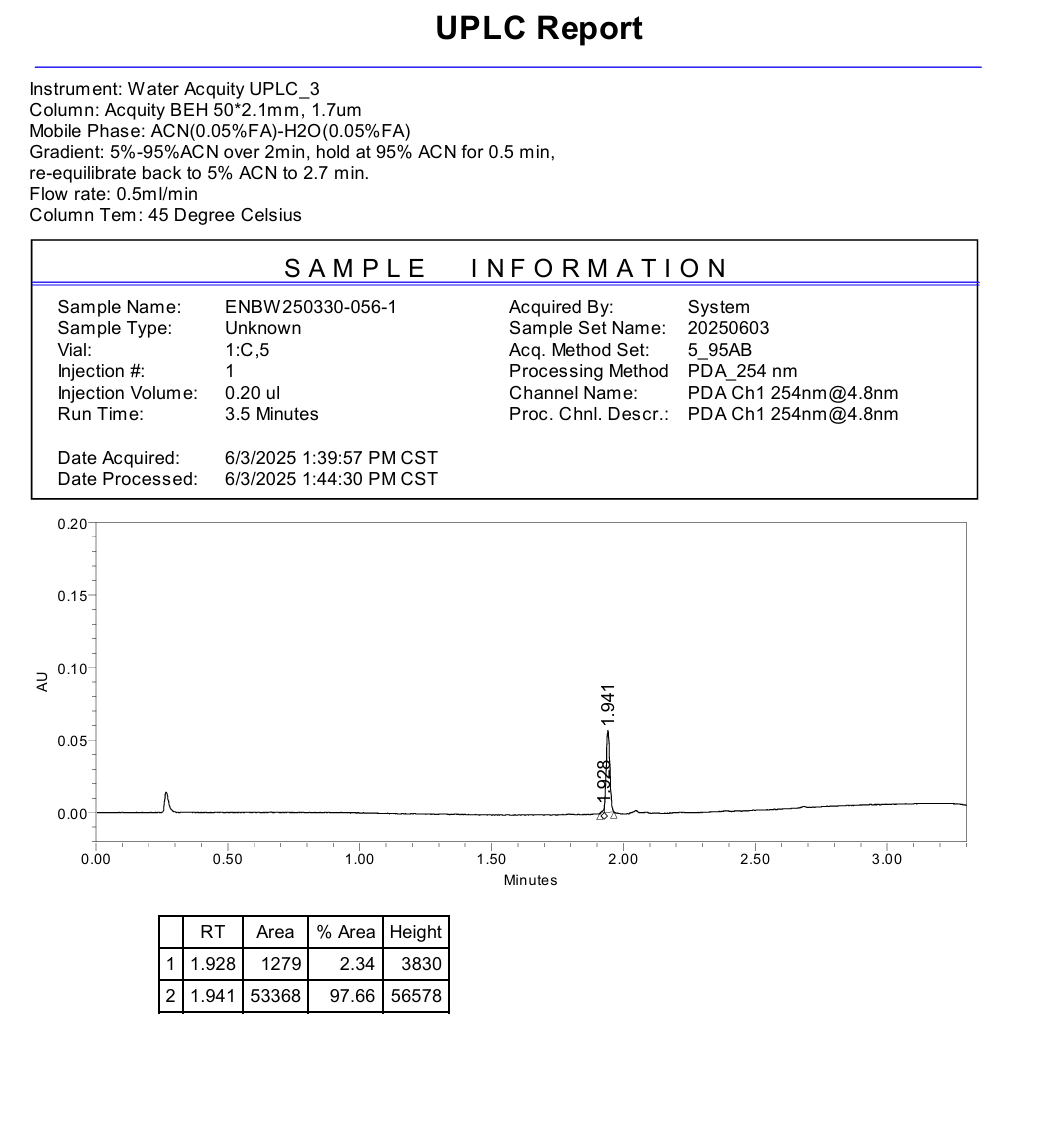

**Fig. S20.** UPLC purity report for **Cpd B**.

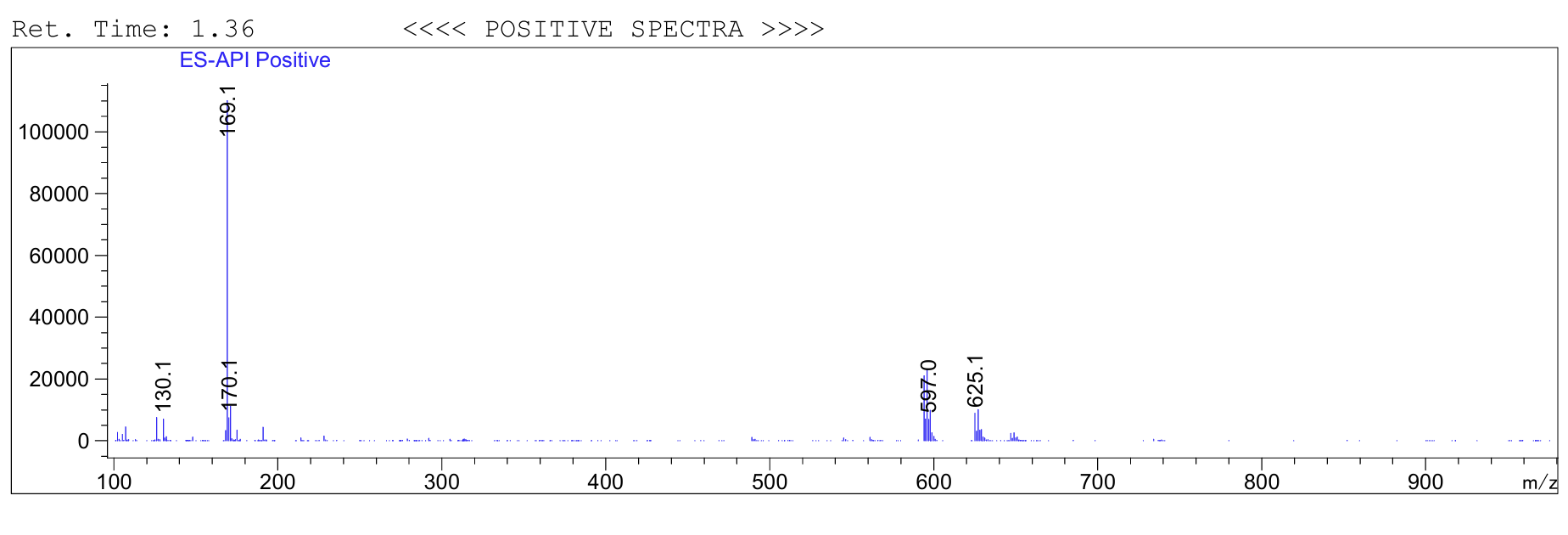

**Fig. S21.**  LCMS for **Cpd B**.

**Crystallographic data collection, refinement and model validation**

**Table S1.** Data collection and refinement statistics. Statistics for the highest resolution shell are shown in parentheses.

| **Data collection** | **CHI3L1–CHI-DEL (PDB 10JV)** |
| --- | --- |
| Number of crystals used | 1 |
| Space group | *P2₁2₁2₁* |
| Cell parameters, a, b, c (Å) | 108.956, 122.309, 136.304 |
| Wavelength (Å) | 0.97934 |
| Unique reflections | 45,095 (3,174) |
| Resolution (Å) | 34.58–2.81 (2.88–2.81) |
| *R*merge (%) | 26.2 |
| Mean *I/σ(I)* | 5.9 (2.22) |
| Completeness (%) | 99.9 (98.0) |
| **Refinement** | |
| Resolution (Å) | 34.58–2.81 |
| Reflections used (test set) | 44,815 (1,997) |
| *R*work/*R*free (%) | 21.5/25.3 |
| **Number of non-hydrogen atoms** | |
| Protein | 11,327 |
| CHI-DEL | 144 |
| NAG and glycerol | 118 |
| Water | 75 |
| **Average B factors (Å²)** | |
| Protein | 52.8 |
| CHI-DEL | 43.5 |
| NAG and glycerol | 80.0 |
| Water | 38.6 |
| **RMSD from ideal geometry** | |
| Bond lengths (Å) | 0.003 |
| Bond angles (°) | 0.679 |
| **Ramachandran statistics** | |
| Favored regions (%) | 97.63 |
| Allowed regions (%) | 2.37 |
| Outliers (%) | 0.00 |
| **Model validation** | |
| Rotamer outliers (%) | 0.42 |
| Clashscore | 6.0 |

**Table S2.** In vitro PK/ADME profile for **CHI-DEL**.

| **PK/ADME parameter** | **CHI-DEL value** |
| --- | --- |
| LogD7.4 | 2.63 |
| Kinetic solubility in PBS, 1% DMSO | 128 µM |
| PAMPA Papp | 5.2 × 10⁻⁶ cm/s |
| MDCK-MDR1 Papp A→B | 7.9 × 10⁻⁶ cm/s |
| MDCK-MDR1 Papp B→A | 10.1 × 10⁻⁶ cm/s |
| Efflux ratio | 1.28 |
| Efflux ratio + elacridar | 1.10 |
| Mouse microsomal t_½_ | 96 ± 8 min |
| Mouse CL_int_ | 19.4 ± 1.8 µL/min/mg |
| Human microsomal t_½_ | 121 ± 11 min |
| Human CL_int_ | 14.8 ± 1.5 µL/min/mg |
| Mouse plasma t_½_ | >180 min |
| Human plasma t_½_ | >180 min |
| Human plasma protein binding | 92.8 ± 0.7% |
| hERG IC_50_ | >50 µM |
| CYP3A4 inhibition at 10 µM | 7.1% |
| CYP2D6 inhibition at 10 µM | 3.9% |
| CYP2C9 inhibition at 10 µM | 5.2% |
| CYP2C19 inhibition at 10 µM | 4.6% |
